# *bofasa* enables high-quality orthology inference across diverse multi-species bacterial genome sets

**DOI:** 10.64898/2026.11.11.765980

**Authors:** Rauf A. Salamzade, Eitan Yaffe, Aamuktha Kottapalli, Lindsay R. Kalan, David A. Relman

## Abstract

Bacterial species can be separated by billions of years of evolution. However, studies of such ancient relationships often rely on methods developed for eukaryotes, which are less diverse. Such methods can regard instances of horizontal gene transfer as contamination, rely heavily on a species phylogeny when partitioning paralogs, and fail to account for the multi-domain architectures of some genes. Here, we introduce, showcase, and benchmark ***bofasa***, a software for high-quality and automated inference of ortholog groups across distantly related bacterial species. bofasa employs a recursive algorithm to split coarse homolog groups into paralogous subsets and first establishes domain-level ortholog groups prior to assessing evolutionary relationships between entire proteins. In comparison to other methods commonly used for multi-species bacterial comparative genomics, bofasa is substantially more conservative in predicting orthology yet can often recover the largest single-copy core genome. Beyond orthology inference, the software also features several additional utility functionalities, including assessment of which ortholog groups are commonly found on mobile genetic elements or possess variable genomic contexts.

## INTRODUCTION

Some molecular estimates suggest the earliest forms of bacteria emerged 3.5 billion years ago^1^. Since this time, bacteria have diverged into over 150 different phyla^2,3^ representing a vast assortment of species. Their genes have also diverged, both in sequence and function. Comparative genomics is a powerful array of methods that mostly rely on using sequence conservation to infer genes that encode for equivalent functions across different taxonomic contexts. Yet, algorithms for examining such gene-to-gene relationships across multiple species have primarily been developed with a focus on multicellular eukaryotic genomes that originated and diversified relatively recently in comparison to the ancient domain of Bacteria. The architecture and evolution of eukaryotic genomes is also fundamentally different than that of bacteria.

Although bacteria are the most diverged domain of life, orthology prediction software designed specifically for their analysis have largely been limited to resolving the pangenome of a single species. Indeed, in this space, highly efficient and accurate tools have already been developed^4–6^ that typically assess the surrounding genomic contexts for ortholog groups to resolve paralogous relationships and leverage high-throughput protein clustering approaches^7,8^. However, for multi-species bacterial comparative genomics, where requirements for syntenic conservation might reduce detection of functionally equivalent ortholog groups, options remain limited^9,10^.

The need for refined tools is imperative because over eons, as bacteria have speciated, their genomes have undergone complex evolutionary events extending beyond mutations, such as gene duplications and horizontal gene transfers (HGT). This has led to intricate evolutionary relationships between ancestrally related genes across different species. While terms such as *orthologs*, *paralogs*, and *xenologs* are clearly defined, accurately discerning such relationships between genes can be difficult. Challenges are especially evident when investigating orthology between distantly related genomes as well as when genes are large and are composed of a mosaic of sequences derived from multiple ancestral sources^11^. *If only a sub-region between two genes is ancestrally related but the genes harbor highly distinct domain architectures, should we assume their protein products are orthologous and perform the same function in their respective cellular contexts?*

As with many areas of biology, this question does not have a clear answer. Some recent methods have begun accounting for complex domain architecture^12,13^ when determining *ortholog groups*, clusters of genes that derive from a common ancestral form; however, many software packages for orthology inference do not do this. Established methods for multi-species orthology inference that ignore domain composition differences thus tend to *over-*cluster proteins that have different functions together^14^. While *under*-clustering of ortholog groups is also problematic and could lead researchers to falsely infer that certain species lack specific functions, having ortholog groups that confidently correspond to equivalent cellular functions should be prioritized.

Here, we introduce a new bioinformatics software tool, *<u>b</u>acterial <u>o</u>rthology finding <u>a</u>nd <u>s</u>yntenic <u>a</u>nalysis* (*bofasa*), enabling automated and high accuracy orthology prediction with methods specifically designed for comparative investigations of diverse, multi-species bacterial genome sets. We showcase its performance in relation to other tools for multi-species orthology prediction with a focus on: (*i*) simulated and real bacterial genome sets, (*ii*) integrases across the genus of *Staphylococcus*, (*iii*) large, multi-domain biosynthetic genes from the phylum *Actinomycetota*, and (*iv*) incomplete genomes representative of draft-quality metagenome-assembled genomes (MAGs).

## RESULTS

### *bofasa* and other software for orthology inference on multi-species bacterial genome sets

The *bofasa* software package features two main executable programs, *bofasa prep* and *bofasa run* (**Fig. 1; Supplementary Figure 1**). The former can accept a directory of bacterial genomes or, alternatively, directories of gene annotations by Prokka^15^ or Bakta^16^, commonly used software to predict bacterial gene coordinates and functions. In addition to vetting the input genomic data for expected formatting, *bofasa prep* will “chop-up” proteins into smaller stretches of sequences that correspond to domain and inter-domain regions. Further, to enable downstream assessment of which ortholog groups are associated with mobile-genetic-elements (MGEs), *bofasa prep* has an option to detect phages and plasmids within the input genomes using geNomad^17^. It also annotates proteins homologous to insertion element (IS)-associated proteins^18^ since they are often found in multiple copies. Another parameter of *bofasa prep* further controls whether such autonomous MGEs should be extracted from their genomic contexts and treated as independent entities during the process of orthology prediction. This enables proper handling of paralogous genes that might exist in multiple copies within genomes across different MGE contexts; however, the option should only be used if genomes are complete and MGE instances are not fragmented due to assembly issues.

**Figure 1:**
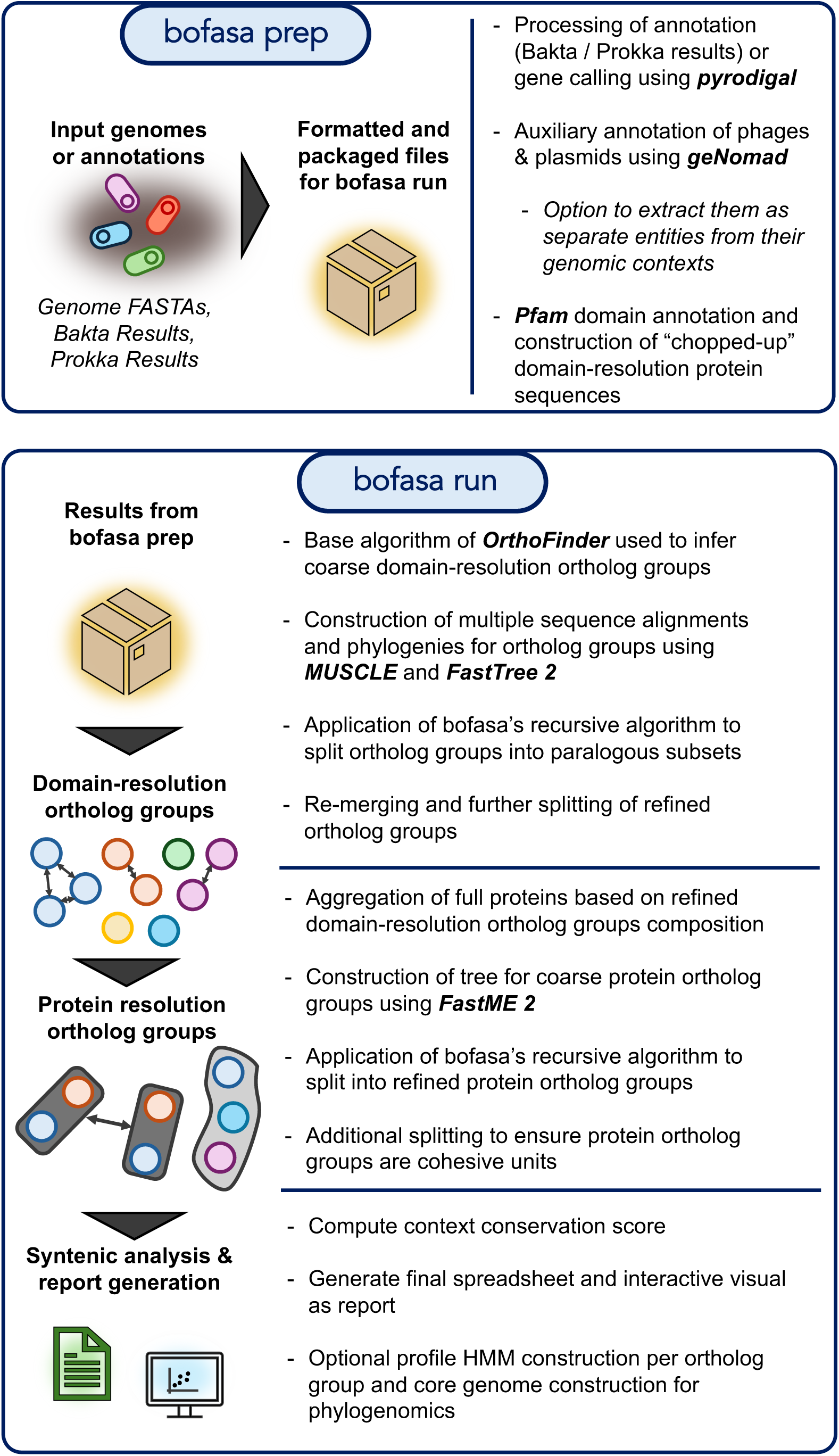
Overview of bofasa prep and run. An overview of the major steps of bofasa prep and run are provided. Key dependency software used in bofasa are highlighted in bold.

The program *bofasa run* then processes formatted data from *bofasa prep* to perform domain-resolution orthology and, subsequently, protein-resolution orthology inference. For determining coarse homology groups at the domain-resolution, *bofasa* applies OrthoFinder’s original algorithm^19^ to adjust scores for sequence length differences prior to clustering via MCL^20^. Afterwards, *bofasa*’s recursive, phylogeny-based algorithm is applied to delineate more resolute domain-level ortholog groups where appropriate (**Fig. 2**). This algorithm essentially aims to identify phylogenetic clades which minimize the number of paralogs; however, additional post-processing is performed to further split or merge back the domain-resolution ortholog groups. Once more granular domain-resolution ortholog groups are finalized, similarities between full pairs of proteins based on domain composition are assessed. Domain overlap between pairs of proteins is used to establish coarse protein families and similarly used to construct distance-based trees with FastME^21^ that are then further split via the same recursive algorithm bofasa uses to refine domain-resolution ortholog groups. Similar to refinement of domain-resolution ortholog groups, additional partitioning of full protein ortholog groups is performed. Finally, *bofasa* calculates a context conservation score for each protein ortholog group and prepares some auxiliary results files, such as an interactive HTML report showing how ortholog groups are distributed according to their context conservation score and prevalence. Users can also request for *bofasa* to determine consensus sequences for ortholog groups as well as construct alignments for near-core proteins for use in multi-partition phylogenomic modelling^22^.

**Figure 2:**
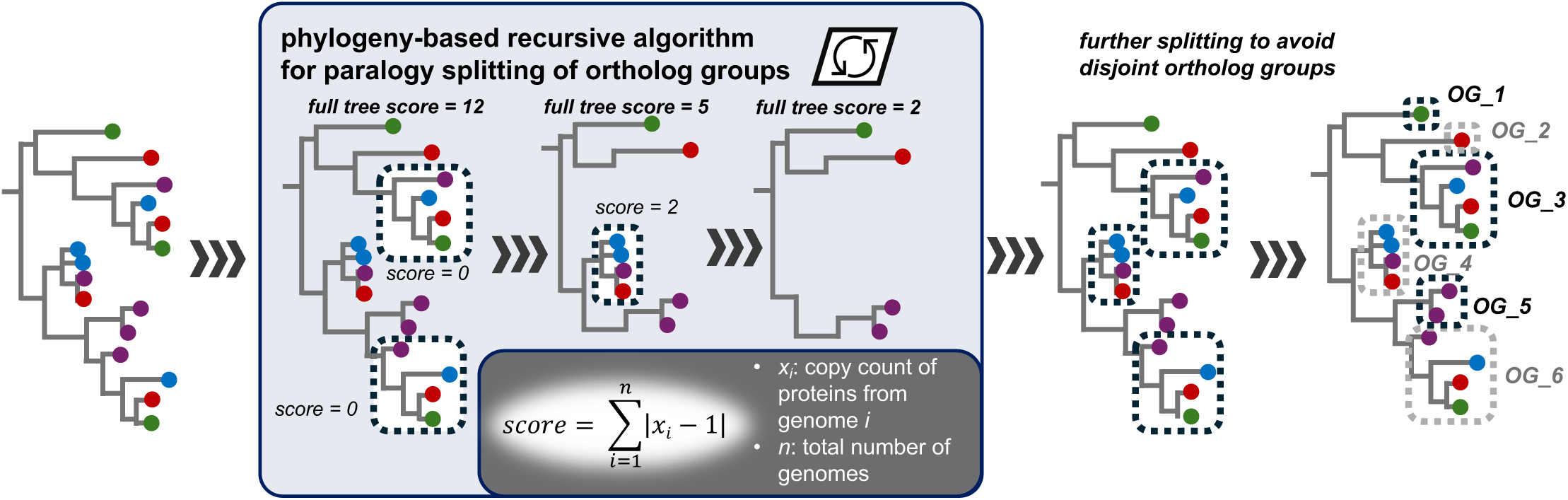
Phylogenetic refinement of ortholog groups in bofasa. Overview of bofasa’s tree-splitting recursive algorithm to split paralogous subsets belonging to a broader homolog group is demonstrated on a simple example tree.

Here, we compare and showcase the performance of *bofasa* to four other methods for multi-species orthology prediction: OrthoFinder^23^, SonicParanoid2^13^, SCARAP^10^, and PIRATE^24^. OrthoFinder is a popular software for orthology inference that has been found to outcompete similar software on primarily eukaryotic testing sets^25^. However, it is mainly designed for eukaryotes, can regard possible instances of HGT as contamination, and does not distinctly handle multi-domain proteins. SonicParanoid2 is another method for orthology prediction which uses various heuristics to improve runtime in comparison to OrthoFinder. It can also use domain architecture to aid orthology inference between multi-domain proteins. Then, PIRATE and SCARAP are orthology software designed for bacterial application, each employing distinct strategies. Finally, for some showcases, we also annotate genomes using the KOfam^26^ database of profile HMMs for KEGG ortholog groups (KOs) to compare how *de novo* orthology prediction software compare to approaches where proteins are assigned to pre-established ortholog groups.

### *bofasa* can recover the largest single-copy core genome while also avoiding over-clustering

We first assessed the accuracy of bofasa and comparator methods on simulated genomes generated using SimPan^27,28^ (**Fig. 3a**). These genomes were derived from an artificial bacteria-like ancestral genome which was simulated to evolve into 20 extant descendants. These 20 extant descendant genomes used for benchmarking were requested to exhibit an average nucleotide identity of 80% between orthologous genes and to depict sequence diversity typically observed across a bacterial genus. Pairs of proteins belonging to common ortholog groups were tabulated for each software and compared to the set of true pairs of orthologs tracked during the process of simulating genome evolution. Methods designed for broad application to eukaryotic and bacterial genomic datasets, such as OrthoFinder and SonicParanoid, typically resulted in over-clustering ortholog groups and an increase in false positive orthology predictions in comparison to methods developed specifically for bacteria. On the other hand, bacteria-specific methods, such as PIRATE and bofasa, tended to miss a greater number of true orthology predictions to avoid making false predictions. SCARAP, a recently developed orthology inference software for application to multi-species bacterial genome sets, defied this trend and, similar to OrthoFinder and SonicParanoid, prioritized minimization of missed orthology relations. Overall, bofasa performed similar to PIRATE but inferred 329 fewer false negative, but also 165 fewer true positive orthologous predictions when both programs were run using default configurations. These differences further increased when bofasa was run with “--random-seed” set to 1000, which requests bofasa to perform a more exhaustive and accurate assessment for splitting up coarse domain ortholog groups into more resolute units. More specifically, this option allows for testing up to 1000 alternate rootings of phylogenies for domain-resolution ortholog groups instead of just applying midpoint rooting. Manual phylogenetic examination of ortholog groups which were affected by over or under-clustering clarified that bofasa’s algorithms worked as expected and such cases were simply difficult to resolve due to multiple options for partitioning that were similarly likely (**Supplementary Figure 2**).

**Figure 3:**
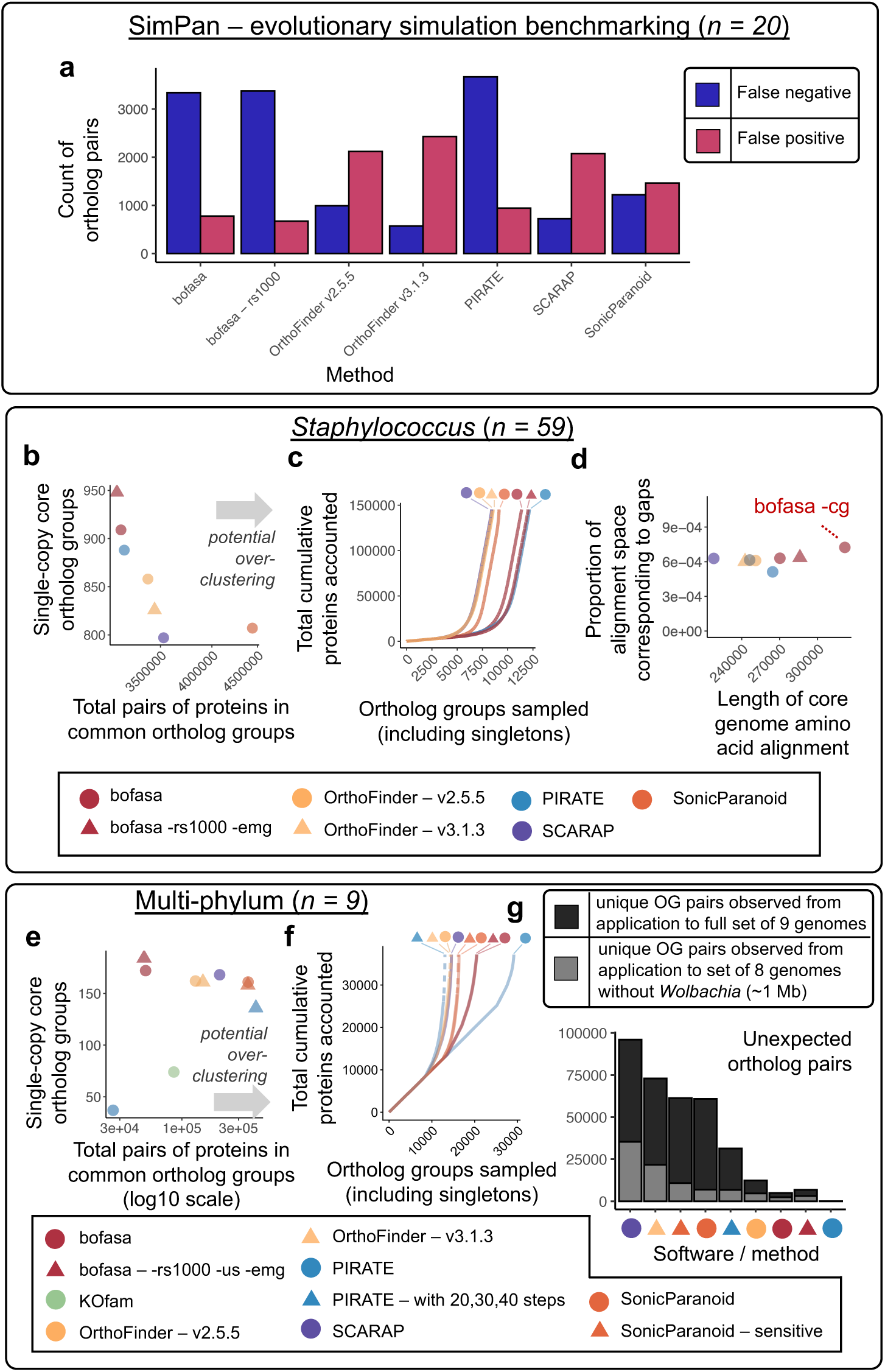
bofasa recovers large single-copy core gene sets while avoiding over-prediction of orthology. (**a**) The number of protein pairs representing false negative (true orthologs that were missed) and false positive (false orthologs that are predicted) orthology predictions are shown for different software through application to a simulated pangenome of a bacterial genus using SimPan. (**b**) The total count of protein pairs co-found in ortholog groups (x-axis) and the number of ortholog groups which represent the single-copy-core genome of the input set of genomes (y-axis) is shown for methods when applied on 59 staphylococcal genomes representative of distinct species. (**c**) The cumulative accumulation of proteins across ortholog groups in order from smallest (singleton) to largest ortholog groups and (**d**) the length and quality (frequency of gaps) of core genome alignments for the same application to staphylococcal genomes. Core genome alignments were trimmed for sites that were largely non-ambiguous. Note, the point labeled as “bofasa -cg” uses the 95% near single-copy core domain ortholog group set whereas the others use strict single-copy core proteins identified by the respective methods. In addition, other differences are present for how the core genome is computed for this point compared to the others, please see the Methods for details. Analogous plots to those in panels (**b**) and (**c**) are shown in (**e**) and (**f**) for a set of nine highly-diverse genomes spanning three separate phyla. (**g**) One of the nine genomes, *Wolbachia*, was excluded and orthology was recomputed *ab initio* using each method. The number of orthologous protein pairs that were distinctly observed were assessed between analogous runs of the same method on the two related datasets. Orthologous protein pairs involving proteins from *Wolbachia* were disregarded.

In order to further assess the performance of bofasa in comparison to established methods, we next performed orthology prediction on a set of 59 complete *Staphylococcus* genomes. Each of the 59 genomes represented distinct species within the diverse genus. Unlike for the SimPan dataset, true orthologous relationships are unavailable for this dataset. We thus compared the performance of different methods using other metrics. First, we assessed their ability to recover a large set of single-copy core ortholog groups while limiting the number of protein pairs found to belong to common ortholog groups (**Fig. 3b**). When run using default configurations, bofasa found 21 more single-copy core ortholog groups than PIRATE while reporting fewer pairs of proteins as belonging to common ortholog groups. It further resolved even more single-copy core ortholog groups when the “--random-seed” option was increased to 1000. In contrast to bofasa and PIRATE, the remaining methods inferred many more pairs of orthologous proteins. This trend is due to these methods producing more ortholog groups that are large -circumscribing multiple paralogous proteins from individual genomes - and is indicative that they over-cluster when applied to bacterial genomes, something that has been previously reported^14^ (**Supplementary Figure 3**). However, it is worth noting that bofasa and PIRATE were unable to group 3.67-4.19% of proteins which were retained as singleton entities (**Fig. 3c**). While this is almost double the percentage observed for other methods, 2.12-3.21%, it is still substantially lower than the percentage of singleton proteins that result from simply binning proteins into pre-computed and curated ortholog groups such as KOfam, which is 73.57%.

As expected, the methods that identified greater numbers of single-copy core ortholog groups also led to larger core genome alignments (**Fig. 3d**). Using the “--core-genome” option of bofasa automates extraction and alignment of a core genome for phylogenomic modeling. This flag requests bofasa to identify near single-copy core (95%) domain-resolution ortholog groups. Domain-resolution ortholog groups are used because such structurally integral regions along proteins are more likely to be evolving under selective pressure as opposed to neutral evolution^29^. Critically, it allows for the incorporation of ortholog groups where a small outlier set of samples might have paralogous instances, ignoring them as if they lacked any instance of the ortholog group. In addition, ortholog groups associated with insertion sequences, phages, or plasmids are automatically discarded from being part of the core genome. Ultimately, this mode enables extraction of a large core genome mostly reflective of sites within coding domains that are evolving vertically.

In addition, we assessed the performance of orthology inference software on a dataset representing phylogenetic diversity beyond the intended scope of the majority of tools. Prediction software were applied on a set of nine extremely diverse genomes, corresponding to genera from three separate phyla: Actinomycetota, Bacillota, and Pseudomonadota. The largest and second largest set of single-copy core ortholog groups identified were by bofasa run with “--random-seed” set to 1000 and bofasa run using default configurations, respectively (**Fig. 3e**). The only method which predicted fewer pairs of proteins as belonging to common ortholog groups was PIRATE; however, it was also unable to recover many single-copy core ortholog groups because its default sequence identity cutoffs are too high for application on such diverse genomic datasets (**Fig. 3f**). In contrast, the default configurations of bofasa were optimized for robust clustering in application to both this dataset and the simulated dataset generated by SimPan, which is substantially less diverse (**Supplementary Figure 4**). Besides bofasa, annotation with KOfams was similarly able to largely avoid over-prediction of orthology and still recover a large single-copy core gene set. However, KOfams’ performance likely owes to the limited number of genomes in the dataset, many of which correspond to model organisms where genes have been more extensively functionally characterized. Amongst the remaining methods, SCARAP was able to recover the second largest set of single-copy core ortholog groups after bofasa but predicted substantially more orthologous relationships.

To further assess the quality of predictions by different methods on the dataset of nine diverse genomes, we excluded one of them, *Wolbachia*, the smallest genome, and reperformed orthology inference. This enabled comparison of orthology inference results when one genome is included versus when it is excluded. Because the remainder of the eight genomes are equivalent between runs being compared, we expect that ideal orthology inference programs should identify the same pairs of proteins between those genomes as orthologs. In accordance with this expectation, bofasa, run with standard and adjusted parameter settings resulted in largely equivalent orthology predictions regardless of whether *Wolbachia* was included or not (**Fig. 3g**). While running PIRATE with default parameters produced the least discordance for pairs of proteins regarded as orthologs between comparing runs, this configuration was also unable to recover many single-copy core ortholog groups. In contrast, some methods which were able to recover a large single-copy core set of ortholog groups produced a large number of discordant ortholog predictions, like SCARAP. Generally, the inclusion of *Wolbachia* resulted in more orthologs being predicted between other genomes. This is potentially because sensitivity for the detection of remote orthologs improves when additional proteins - that can serve as steppingstones - are included.

### *bofasa* orthology analysis can resolve complex evolutionary scenarios involving mobile genetic elements

Some bacterial genomes harbor autonomous entities, commonly referred to as mobile genetic elements (MGEs), that are able to laterally transfer between cells of the same or different species^30^. Two major types of MGEs include phages and plasmids, which can be annotated along genomes using geNomad^17^. In addition to integrating geNomad for determining whether any ortholog groups largely correspond to MGE-associated proteins, bofasa also implements the option “--extract-mge-genomes” to support automated extraction of phages and plasmids to be considered as separate genomic entities during orthology inference. To showcase this option and bofasa’s calculation of ortholog group specific context conservation scores, we further investigated the set of 59 *Staphylococcus* genomes representative of distinct species.

After running bofasa on the input set of genomes, the relationship between ortholog group context conservation scores and frequencies was assessed (**Fig. 4a**). In comparison to other software for orthology inference and bofasa run with default parameters, bofasa run with the flag “--extract-mge-genomes” resulted in the unique identification of a large ortholog group, OG001033, that corresponds to predicted phage-associated integrases (**Supplementary Figure 5a**). Integrases are enzymes that enable MGEs to integrate within genomes and around 67% of the instances of this ortholog group were found within regions annotated as phages by geNomad^31^. To further understand the evolution of the OG001033 integrase family, we first identified additional ortholog groups containing protein domains that have been associated with integrases^32^ (**Fig. 4b**). Surprisingly, the context conservation for most integrase-related ortholog groups fell within expected ranges when accounting for ortholog group frequency. Some ortholog groups that were highlighted even depicted high degrees of context conservation and further annotation revealed that they correspond to house-keeping genes. For instance, one ortholog group mapped to the gene *xerC*, which encodes a tyrosine recombinase shown to be essential for proper chromosomal segregation but can also be co-opted by phages^33,34^. In addition, OG001033, was found to harbor a handful of proteins that featured no integrase-associated domain. Depending on parameters used, between 66.7% and 72.7% of such putative *integrases by association* identified by bofasa were regarded as likely integrases based on structure predictions^35,36^. These rates are higher than for other methods besides SonicParanoid (**Supplementary Figure 5b**).

**Figure 4:**
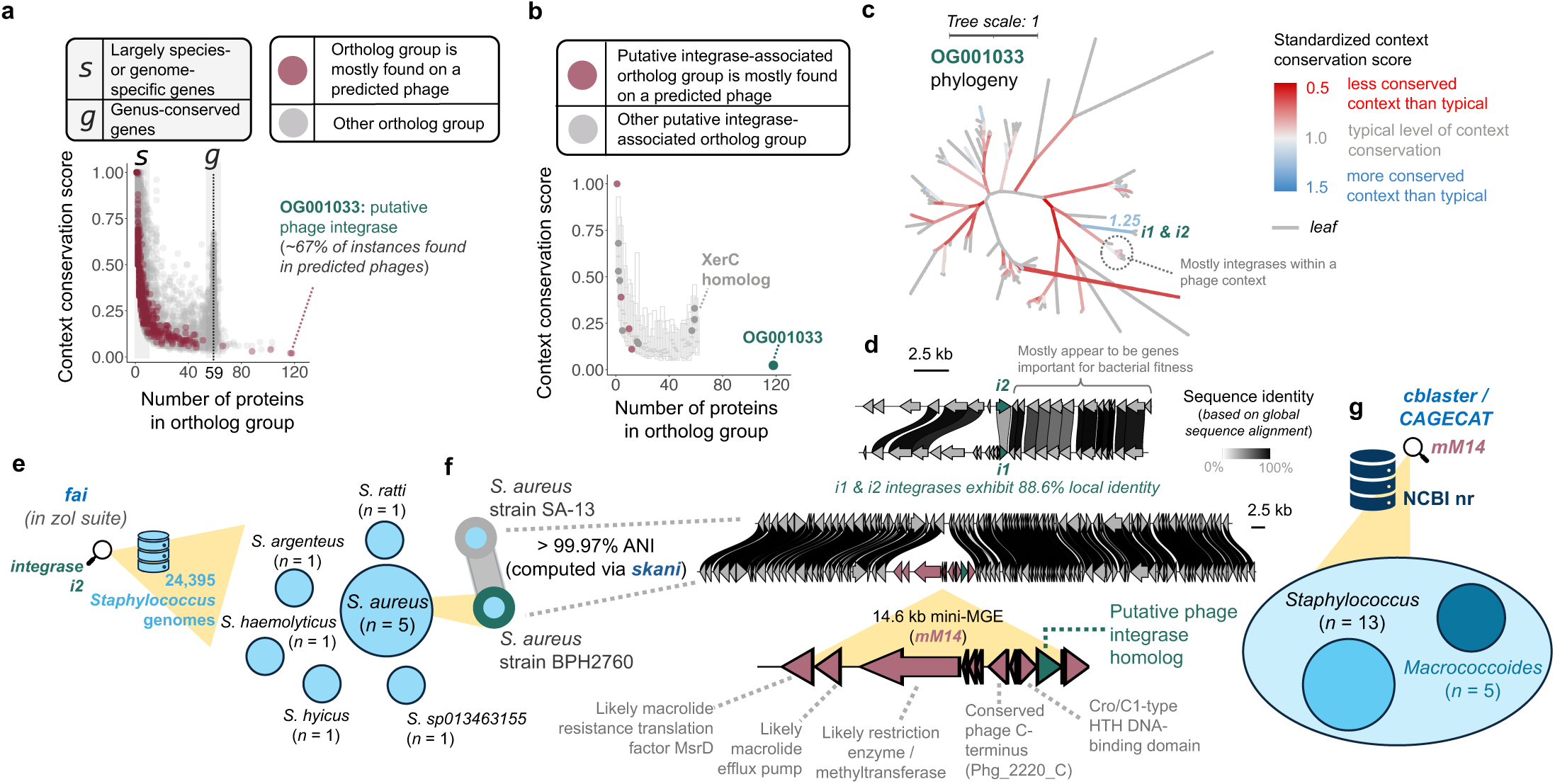
bofasa-based investigations of the *Staphylococcus* genus are congruent with functional expectations. (**a**) The relationship between the number of proteins found in ortholog groups and their context conservation score (ccs) is shown. If ≥50% of instances are found on a predicted phage, ortholog groups are shown in brick red instead of grey. The approximate locations of ccs peaks for species or genome specific [*s*] and genus-wide conserved [*g*] genes. (**b**) An alternate view of (**a**), where for each protein count bin where there are 20 or more ortholog groups, a distribution is determined to construct skeletal box plots where the mid-line corresponds to the median ccs, the bottom-boundary represents the 10^th^ percentile value, and the upper-boundary represents the 90^th^ percentile value. Ortholog groups containing integrase-associated domains are highlighted in brick red or grey. (**c**) A phylogeny of the 118 proteins belonging to OG001033. The coloring of the branches coincide to the standardized ccs, which is the ccs divided by the median or expected ccs given the number of proteins regarded as children of the inner-tree branch. (**d**) A clinker-based visualization of the contexts of integrases *i1* and *i2*. (**e**) A fai-based search for integrase *i2* in single-query mode revealed that only 10 genomes harbor a variant that exhibits ≥88.6% identity. (**f**) Using skani, the strain SA-13 was identified as closely related and used to determine the boundaries of the mini-MGE, *mM14*, harboring the *i2* homolog. (**g**) The cblaster software suite was used to remotely search for additional instances of *mM14* in NCBI’s nr database, revealing five instances within genomes of the closely related genus *Macrococcoides*.

While context conservation scores are computed by bofasa for each protein ortholog group, they can also be computed for any subset of proteins. After constructing a phylogeny of proteins belonging to the ortholog group OG001033, we calculated the context conservation score for each branch leading to an internal node and standardized the scores by the median context conservation score expected for the number of proteins being considered (**Fig. 4c**). This revealed that most integrase enzymes within this ortholog group have largely different contexts. Even closely related proteins, despite being found integrated within the genomes of different staphylococcal species, have ±10 kb contexts that exhibit less conservation than would be expected. This suggests: (i) that integrases might frequently change contexts and/or (ii) that integrated MGEs, like prophages, rapidly evolve relative to other regions of bacterial genomes. In contrast to most branches, one phylogenetic partition corresponded to a pair of integrases, *i1* and *i2*, that share a more conserved context than expected (**Fig. 4d**). The surroundings of both integrases included two highly conserved genes encoding for ribosomal proteins, *rpsI* and *rplM* (aa percent identities: 94.6-95.2%), and a set of three other conserved proteins (aa percent identities: 88.1-96.1%)^37^. While phages can carry ribosomal proteins and phylogenetic neighbors of *i1* and *i2* were mostly found within phage contexts, comparative genomics suggested that these ribosomal proteins are likely part of the staphylococcal chromosome similar to the genes that lie upstream of them^38,39^ (**Supplementary Figure 6**).

To further investigate this integrase, we holistically searched for *i2* across 24,395 *Staphylococcus* genomes^40^ (**Fig. 4e**). Only ten integrases were found where the amino acid identity based on local alignment was greater than or equal to 88.6%, which is what was observed between *i2* and *i1*. Five of these instances were found in *Staphylococcus aureus* genomes. We leveraged the abundance of genomes for *S. aureus,* a well-studied and well-sequenced pathogen, to identify a genome lacking the integrase but possessing a high sequence similarity to a genome with it^41^. Comparative analysis between these two genomes revealed an evident mobile genetic element, *mM14*, that is approximately 14.6 kb and harbors genes putatively related to macrolide-resistance^37,40^ (**Fig. 4f**). Searching for instances of *mM14* in NCBI’s nr database revealed several highly homologous instances within *Staphylococcus* and *Macrococcoides* genomes^42^ (**Fig. 4g**). While some genes shared between these *mM14*-like elements exhibited high-identity, others were variably carried and MGE-specific (**Supplementary Figure 6**).

Ultimately, this showcase highlights how bofasa can be used to determine more comprehensive ortholog groups associated with MGEs that other methods would partition based on genomic context. By further using scalable tools for targeted detection, we were able to identify additional instances of a sub-variant of an integrase family. Our analysis suggests that *i1* and *i2*, each found in a rare staphylococcal species which only has one genome available, are diverged instances of an integrase found in MGEs belonging to *S. aureus*, where it can be co-located with potential macrolide resistance genes. The integrase *i1* further appears truncated, indicating that it might not be functional and a remnant of a decaying MGE.

### *bofasa* alleviates over-clustering of multi-domain proteins and can aid inference on their cellular function

Some proteins, for instance, large non-ribosomal peptide synthetases (NRPSs), feature complex multi-domain architectures^43^. NRPSs are particularly prevalent in *Actinomycetes*, a class within the phylum *Actinomycetota*, known for their production of complex specialized metabolites with various functional roles^44–46^. We thus further compared bofasa to comparator orthology prediction or annotation methods through applications to 56 *Actinomycetota* genomes, each representative of a distinct genus and corresponding to a complete assembly of length five megabases or larger. The largest single-copy core gene set was identified by bofasa. SonicParanoid2 was close to finding a similar number of single-copy core gene as bofasa; however, it reported substantially more orthologs, 6.21-6.26X more. Overall, bofasa reported the lowest counts of ortholog predictions, the highest number of single-copy core ortholog groups detected, and the highest average rate of containment by other methods (**Supplementary Figure 7**).

We next annotated the presence of NRPS containing regions^47^ and assessed orthology predictions for NRPSs across the genome set. Most methods inferred a limited number of ortholog groups and over-clustered (**Fig. 5ab; Supplementary Figure 8**). Besides bofasa, all six alternate methods assessed produced large ortholog groups comprising of 300 or more proteins. In contrast, bofasa delineated more resolute ortholog groups, only two of which were comprised of more than ten NRPSs. Both ortholog groups were also found in biosynthetic gene cluster (BGC) regions which were at least partially predicted to underlie NRPS-based synthesis of metallophores and found in highly divergent taxa, spanning three orders within the class *Actinomycetes* (**Fig. 5bc**).

**Figure 5:**
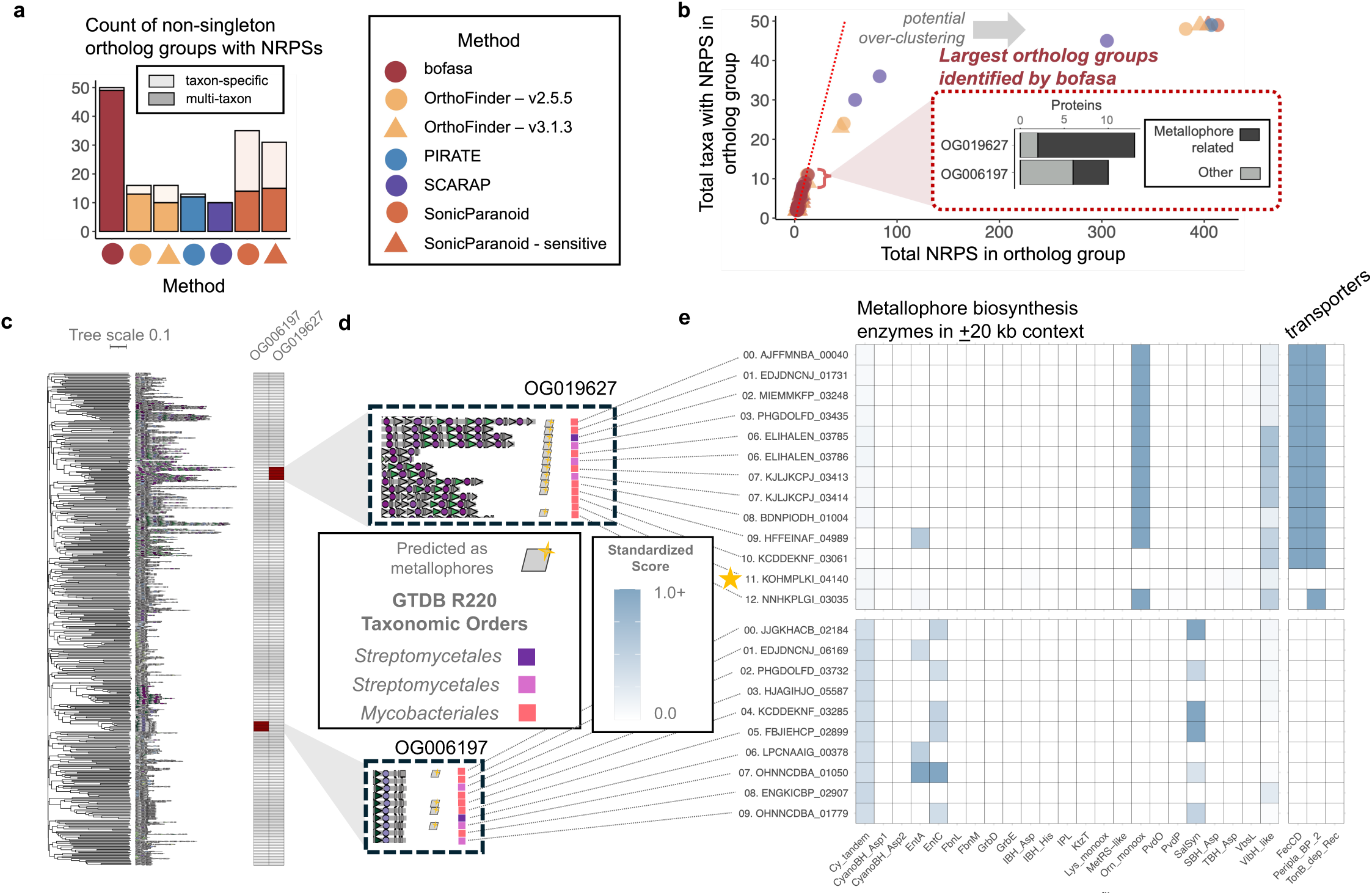
bofasa sheds light on the evolution and context of non-ribosomal peptide synthetases. (**a**) A comparison is provided for the distinct number of non-singleton ortholog groups with non-ribosomal peptide synthetases (NRPSs) with coloring corresponding to whether they are genome/species specific or multi-taxon. (**b**) A scatterplot is shown providing the number of proteins and source genomes non-singleton, NRPS-associated ortholog groups are found in. (**c**) A cosine-distance based tree from domain compositions of NRPSs was constructed using FastME 2. The domain architectures and protein membership for the two largest ortholog groups are shown. (**d**) Additional zoom-ins of the domain architectures for the two ortholog groups are provided alongside markers for whether they were predicted as metallophores. (**e**) The presence of homologs to key metallophore-associated biosynthesis enzymes and transporters cataloged by Reitz et al. 2026 and Crits-Christoph et al. 2021 are shown with coloring corresponding to the standardized score from PyHMMER searches. Values equal to or greater than 1 (blue) indicate sufficient homology to regard them as present. The star indicates the single protein belonging to OG019627 that does not have any clear metallophore-associated enzymes or transporters nearby.

We further investigated the domain architecture and genomic contexts of NRPSs belonging to these two ortholog groups to assess whether they are metallophore specific and were missed due to sequence similarity thresholds (**Fig. 5cde, Supplementary Figure 9**). To do so, we assessed whether proteins in the nearby vicinity of the focal NRPSs matched to key metallophore biosynthesis enzymes^48^. For the larger ortholog group, OG019627, all but two instances were found nearby the metallophore biosynthesis profile for ornithine N-monooxygenase. We also found support that one additional BGC without this profile could also underlie synthesis of a metallophore due to the nearby presence of specific transporter domains to the NRPS^49^. Interestingly, the remaining NRPS exhibited clear orthology to other proteins in this group but lacked such transporters or any enzymes exhibiting high homology to known metallophore biosynthesis profiles. For the smaller ortholog group comprised of ten proteins, only four were predicted as potentially related to metallophores based on default criteria. The contexts for three of the four instances exhibiting greater support for underlying metallophore biosynthesis featured bifunctional salicylate synthases and the other featured an isochorismate synthase and 2,3-dihydro-2,3-dihydroxybenzoate dehydrogenase. There is no clear presence of metallophore-associated transporters or biosynthesis enzymes in the contexts of the other NRPSs; however, all ten instances featured low level homology to domains for tandem heterocyclization (36-43% of the needed score cutoff for detection). Together these results suggest two major possibilities: (1) some of these NRPSs might be nearby novel or diverged metallophore biosynthesis genes, and / or (2) that these orthologous NRPSs can be found within both BGCs related to and unrelated to metallophore biosynthesis.

### *bofasa* can be applied to investigate MAGs from a single microbiome to perform comparative metagenomics

Due to advances in sequencing technologies and bioinformatic methods, the availability of high-quality metagenome assembled genomes (MAGs) is quickly rising. This brings the exciting prospect of performing comparative genomics between MAGs constructed from a single microbiome sample. To showcase bofasa’s utility for such applications, we focused on the hundred most abundant bacterial species found in stool samples from 60 healthy participants and gathered representative complete genome assemblies for them^50^. Similar to other showcases, in comparison to other methods, we found that bofasa achieved a good balance in resolving a large number of single-copy core genes while limiting the total number of orthologs predicted (**Fig. 6a**). bofasa once again produced substantially fewer large ortholog groups comprising of more proteins than the number of genomes investigated. In addition, as demonstrated previously, a taxonomically diverse dataset can result in an increase of false orthology predictions. To further quantify the impact of this, we assessed the ratio of the total number of domains found within ortholog groups to the average number observed for individual member proteins (**Fig. 6b**). Higher values of this ratio indicate ortholog groups that circumscribe proteins with more distinct domain architectures and thus likely a multitude of cellular functions. In addition to emitting a limited number of large ortholog groups, the large ortholog groups that bofasa did produce also had relatively cohesive domain architectures in comparison to alternate methods (**Fig. 6c**).

**Figure 6:**
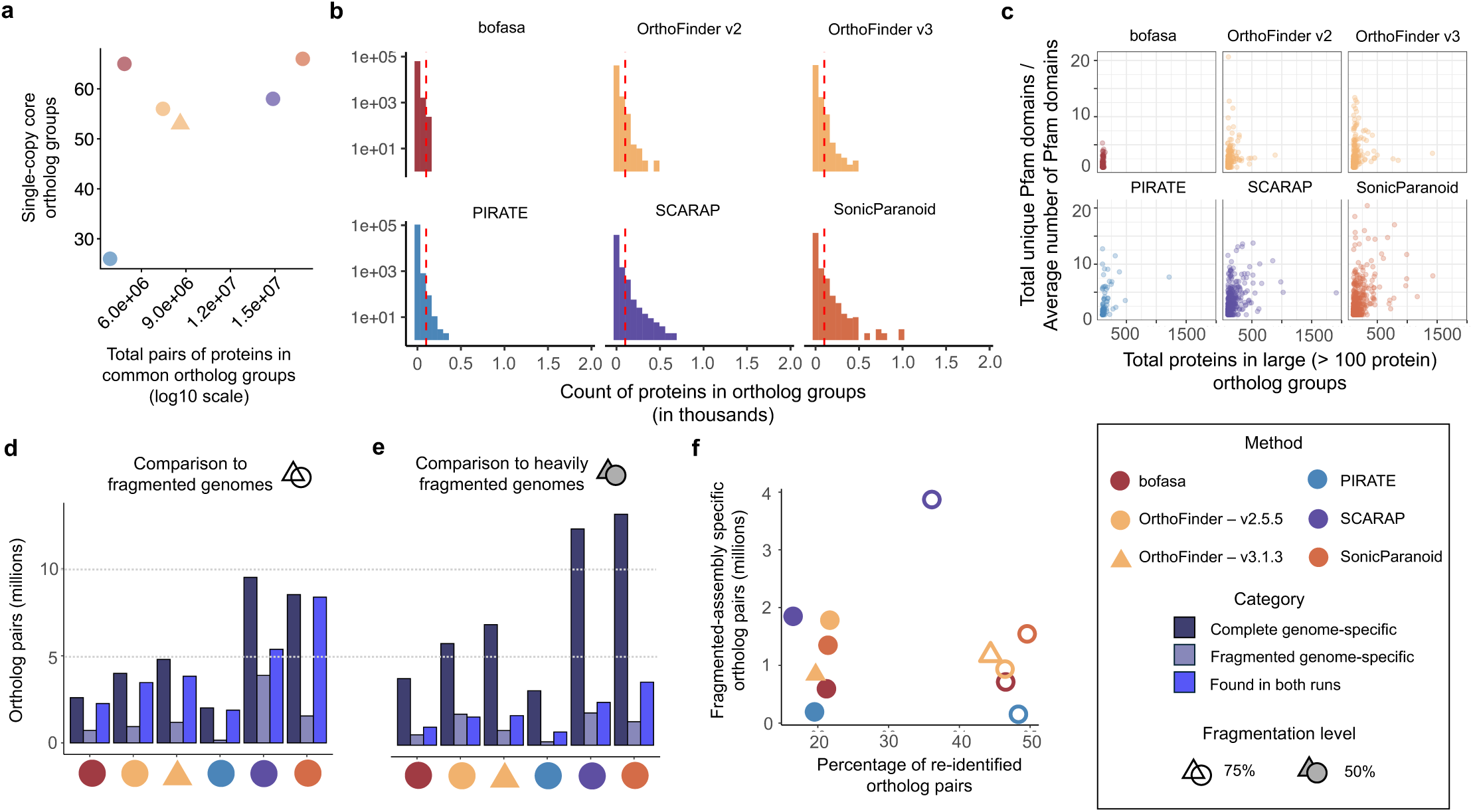
bofasa acts conservatively for orthology inference when applied to a diverse genome set representative of the gut microbiome. (**a**) The total count of protein pairs co-found in ortholog groups (x-axis) and the number of ortholog groups which represent the single-copy-core genome of the input set of genomes (y-axis) is shown for different prediction methods when applied to 100 representative genomes that are abundant in the healthy gut microbiome from re-analysis of Yaffe et al. 2025. (**b**) The size distribution of ortholog groups predicted by different methods. (**c**) The total number of proteins in large ortholog groups, those exceeding the number of source genomes, is shown alongside a statistic corresponding to the total number of distinct Pfam domains standardized by the average number of such Pfam domains. (**de**) Assembly fragmentation was simulated on the set of 100 genomes at two different levels. The categorizations correspond to the number of ortholog pairs that are shared or distinct between equivalent runs by methods on the complete genomes and simulated fragmented genomes. (**f**) The relationship between the percentage of re-identified ortholog pairs from re-running orthology prediction on simulated fragmented genomes and the count of ortholog pairs only identified when orthology is inferred using fragmented genomes (likely false positives) is shown.

While long-read sequencing is resulting in high-quality MAGs, many MAGs, in particular those generated using short-read sequencing, are often of modest contiguity. While we previously assessed how different methods for orthology prediction perform when including and excluding a single genome (**Fig. 3g**), using the set of 100 complete genomes representative of healthy gut microbiomes, we next assessed how genome incompleteness might affect orthology predictions. To do so, we simulated artificially fragmented variants roughly corresponding to 75% and 50% completeness by randomly introducing gaps of varying length. Comparison of orthology predictions revealed that all methods, including bofasa, were vulnerable to missing ortholog pairs when run on incomplete genomes and also culpable of producing orthologous predictions that are absent when the same methods are run on complete genomes (**Fig. 6def**). This likely reflects issues with reciprocal best hits (RBHs) that are imperfect and also downstream inabilities to resolve coarse homolog groups based on RBHs using phylogenetics. PIRATE produced substantially fewer false predictions that were exclusively observed in fragmented genomes compared to other methods. This likely reflects its stringent default percent identity and coverage criteria for defining orthologs. Ultimately, these analyses highlight a need to be cautious when applying bofasa and other orthology inference methods on sets of genomes that are incomplete.

## DISCUSSION

Here, we introduce bofasa, automated software for orthology inference across multi-species bacterial genome sets. In comparison to other methods, bofasa tends to be more conservative in inferring orthology, extracts the largest single-copy core genome, and avoids production of extremely large ortholog groups. However, bofasa is computationally expensive and can sometimes take longer to run and require more memory, albeit not substantially, than comparator methods (**Supplementary Figure 10**). This makes it ideal for deep evolutionary investigations of 200 or fewer representative genomes but less appropriate for comprehensive investigations of hundreds or thousands of bacterial genomes. As we show in the manuscript, interesting proteins highlighted by bofasa can however be further investigated at scale using software such as BLAST^51^, fai^40^, and cblaster^42^ by performing targeted searches that also allow for the detection of more remote homologs.

We further demonstrated how bofasa’s orthology inference is robust to the inclusion and exclusion of genomes in comparison to other methods, in particular those that are designed for scalability. However, bofasa, similar to many other methods for orthology inference, can infer orthologs or fail to split paralogs when applied to partial genomes, such as MAGs generated using short-read sequencing. PIRATE, which applies strict percent identity cutoffs when inferring orthologs, was the best at limiting such false orthology predictions when dealing with draft-quality genome assemblies. As such, we might incorporate options for users to request strict cutoffs to further refine ortholog groups. Although we purposefully avoid leveraging synteny for orthology inference to not confound secondary syntenic analysis, syntenic information can also be used to further refine ortholog groups and we might incorporate its usage as such later on^52,53^. Similarly, we might eventually incorporate structural similarity assessment within bofasa^36,54,55^, which is currently difficult to scale but will likely become more computationally tractable in upcoming years.

The advantages of bofasa for resolving ortholog groups in bacterial genome datasets are demonstrated here through five showcases, including an option for handling orthology prediction for MGEs such as plasmids and phages. In general, bofasa produced more granular ortholog groups with more cohesive domain compositions than other methods as demonstrated by application to elucidate the evolution of NRPSs in the phylum Actinomycetota. For that showcase, bofasa likely avoided over-clustering distinct NRPSs due to its ability to resolve more granular domain-resolution ortholog groups rather than simply rely on domain composition based on annotation of Pfam profiles like SonicParanoid 2. This would align with improved performance achieved from using sub-Pfam profile HMMs by the BiG-SLICE software package specifically designed to cluster related BGCs^56^. Further, by allowing for autonomous MGEs to be treated as distinct genomic entities, bofasa was also able to resolve a large group of integrases that other methods were unable to holistically aggregate.

Finally, in recent years, there have been significant advances in artificial intelligence (AI) models. To construct such models often requires high-quality training data to learn underlying patterns^57^. While not very scalable itself, bofasa can be applied to genomes of diverse genera to produce high-quality orthology predictions. These predictions can then potentially be used to train AI-based heuristics, such as autoencoders^58^, for more scalable orthology inference.

## KEY ACRONYMS AND DEFINITIONS

HGT: horizontal gene transfer
Orthologs: genes arising from a common ancestor but diverging due to speciation
Paralogs: genes diverging due to duplication
Xenologs: related genes via HGT
MGE: mobile genetic element
NRPS: non-ribosomal peptide synthetase
MAG: metagenome assembled genome

## ACKNOWLEDGMENTS

This work was supported by funding from Good Ventures (D.A.R) and the National Institutes of Health (NIAID U19AI142720; L.R.K.). The content is solely the responsibility of the authors and does not necessarily represent the official views of the National Institutes of Health. We thank members of the Relman and Kalan labs for useful feedback on the development of bofasa and application showcases presented in this manuscript.

## METHODS

There are two major commands in the bofasa software package, bofasa prep, used to process and format input genomic data, and bofasa run, which carries out the actual orthology prediction and other auxiliary functions included in the final set of results. Similar to some other methods, for instance Domanoid^12^, SonicParanoid 2^13^, BiG-SLICE^56^, and BiG-SCAPE^59^, a key component of bofasa’s orthology inference is that it accounts for multi-domain proteins and partial orthology.

### Software availability

bofasa (https://github.com/raufs/bofasa) is open-source, provided under the BSL-3 license, developed in Python 3, and available for download on both Bioconda^60^ and BioContainers^61^. For this manuscript, version 1.2.0 of bofasa was used (specifically, commit bef6947, Jan 22, 2026, for all bofasa prep and run analyses and commit 81a3489 for database setup on Aug 9, 2025). Artificial intelligence (AI) models were not used in the initial development of the software but were used to perform supervised re-structuring of the code and implementation of new, secondary features. Additional details on comparative and benchmarking analyses performed for the manuscript can be found at: https://github.com/Kalan-Lab/bofasa_manuscript. Releases of the code repository (starting with v1.2.0) and the repository for manuscript-specific analyses are also stored in immutable forms on Zenodo^62–64^.

### Input data processing using bofasa prep

bofasa prep processes input genomes or annotation directories to prepare files for the primary program, bofasa run. It takes as input either a set of genomes in FASTA format or sample-specific annotation directories from Prokka^15^ or Bakta^16^. Due to intensive computational requirements throughout bofasa, it is by default restricted to processing a maximum of 200 genomes. A minimum of four genomes is also required.

Beyond formatting and optionally designating new and distinct locus tags, bofasa prep further splits coding sequences into domain and inter-domain subsequences based on PyHMMER^65^ (v0.11.1) annotation with the Pfam-A database^66^ (release 37.4), runs geNomad^17^ (v1.8.1) to determine which proteins are on phage or plasmid contexts, and infers homology to insertion elements^18^. For splitting coding sequences, hits are ordered based on the score and then largely non-overlapping segments are selected. By default, only segments of length 20 amino acid residues are considered. If users are working with complete bacterial genomes, an option is also available, “-emg” or “--extract-mge-genomes”, to extract phage and plasmid regions or contigs as separate entities for orthology inference.

### Orthology prediction workflow in bofasa

The bofasa run workflow primarily determines ortholog groups and takes the resulting directory from bofasa prep as input. The first step in bofasa run is to execute OrthoFinder’s base algorithm^19^ (v2.5.5) for determining coarse ortholog groups using split coding sequences corresponding to domain and inter-domain units. We use OrthoFinder 2 instead of OrthoFinder 3 because benchmarking revealed the former tends to favor specificity over sensitivity when determining initial coarse ortholog group clusters. OrthoFinder’s base algorithm uses DIAMOND blastp^67^ (v2.1.10) and importantly performs correction for gene length, which contributes to it being amongst the most accurate methods for orthology inference^25^. If bofasa prep is run using the ‘-emg’ option, then OrthoFinder is run using only proteins from bacterial chromosomes and afterwards the resulting file is modified to include proteins from phages or plasmids based on DIAMOND blastp alignment to chromosomal and other MGE proteins. Next, multiple sequence alignments and phylogeny construction are run for each coarse domain-resolution ortholog group using MUSCLE^68^ (v5.1) in super5 mode and FastTree 2^69^ (v2.1.11), respectively. Prior to phylogeny construction, protein alignments are trimmed using trimAl^70^ (v1.5.0) in strict mode. Phylogeny construction is based on the trimmed alignment if at least ten sites remain after trimming and the full alignment otherwise.

Afterwards, the phylogeny of coarse domain-resolution ortholog groups is processed to split paralogous groups. By default, midpoint rooting is applied and solely assessed, but multiple internal nodes can be requested for rooting via the ‘-rs’ option. For each rooting, the phylogeny is first investigated with bofasa’s recursive algorithm for splitting trees. This algorithm is inspired by but is different from the one used in OrthoSNAP^71^. Briefly, it finds monophyletic clades within rooted trees which have the lowest value of a score. This score is simply: Z |*c* − 1|, where *c* is the copy-count of proteins from a single genome for all genomes in the initial input tree. The score is thus lowest for clades which have a single protein from each genome. The clade or clades, if there is a tie, with the lowest scores are identified and cataloged, then filtered from the tree which is reprocessed through the recursive algorithm. If the lowest score for subclades is equal to or greater than the score for the full tree, then the algorithm terminates and no further splitting is performed. The algorithm also terminates if the branch lengths of the tree are 0.0005 or lower, indicating there is no variation between tree leaves since 0.0005 is a pseudo-value assigned by FastTree 2 to prevent branch lengths of 0. Afterwards, the initial tree is further processed to split phylogenetically disjointed proteins that might commonly belong to the coarse background ortholog group. This can be the case when a discrete paralogous group is identified within a larger clade and the leftover proteins, outside of the split paralogous group, are phylogenetically disjoint. This splitting is performed by first assessing larger subclades with more children to avoid over-splitting. Finally, a reassessment is performed for merging split partitions of an ortholog group using a fixation index. This works by assessing pairs of split ortholog groups that belong to a common larger phylogenetic clade and computing a fixation index: (*π_bet_*_w*een*_ − *π*_w*it*ℎ*in*_)/*π_bet_*_w*een*_, where *π*_w*it*ℎ*in*_ is the mean branch distance between domain-resolution sequences from the same group and *π_bet_*_w*een*_ is the mean branch distance between domain-resolution sequences from complementary groups. By default, groups are merged if the fixation index for them is below some threshold, 0.1. All three steps -the recursive splitting of paralogous groups, the secondary splitting of disjoint proteins, and merging of paralogous groups based on fixation index -are performed for each rooting of a tree.

When run using multiple rooting options via the “-rs” option, two scores are used to determine which partitioning scheme achieved is the best to use. A primary cumulative score is determined for the entire coarse domain-resolution ortholog group as the sum of scores - the same metric used in the recursive algorithm - across split partitions. A secondary cumulative score is also computed as the sum of intra-clade branch lengths. The best partitioning scheme per ortholog group arising from use of different phylogeny rootings are assessed using the primary score. If there is a tie, then the secondary branch-lengths based score is also assessed. For both metrics, lower scores indicate more high-quality partitioning of coarse ortholog groups into refined ortholog groups.

The refined domain-resolution ortholog groups are next used to determine coarse protein-resolution ortholog groups. This is performed through assessing pairs of proteins for common domain-resolution ortholog groups using a Jaccard index. Specifically, pairs of proteins are linked if they exhibit a Jaccard index greater than or equal to the threshold *D*_j_ − (*D*_j_ ∗ *X*), where *D_j_* is by default set to 0.5 and *X* corresponds to the proportion of total genomes which harbor the most common single-copy domain-resolution ortholog group. This supports linking of proteins which feature a defining domain-resolution ortholog group that is single-copy but otherwise requires a reasonable overlap for domain-resolution ortholog groups. Domain-resolution ortholog groups that are largely inter-domain (>80% of instances) are not considered. Linked pairs of proteins are then used to establish coarse protein clusters via the Markov clustering algorithm (MCL)^20^ with the inflation parameter set to 1.2 to achieve coarse clusters. Resulting ortholog groups with a maximum copy-count of two or greater and featuring proteins from two or more genomes are further investigated to potentially split paralogous subsets where appropriate. This is performed through tree inference via the BalME method in FastME (v2.1.6.3)^21^ using cosine distances of domain-resolution ortholog group carriage profiles computed for pairs of proteins. The trees are then midpoint rooted and processed through bofasa’s recursive algorithm for partitioning ortholog group trees as described for resolving domain-resolution ortholog groups. Disjoint protein sequences are further split into discrete ortholog groups as described previously as well; however, no final merging based on branch-lengths is performed. Instead, after further splitting based on phylogenetic structure, each protein within each resolute cluster is assessed for whether it is as or more similar to a protein from another cluster than it is to any protein in its own cluster. Such proteins are disregarded as members of their respective cluster and categorized as singleton ortholog groups. The assessment between proteins is based on a Jaccard index, during which largely inter-domain ortholog groups are regarded for greater resolution.

Ultimately, two tab-separated matrix files are written to the final results subdirectory for the refined, granular domain-resolution and full-protein ortholog groups. In addition to these matrix files, an XLSX spreadsheet and an interactive HTML visual are also produced. The XLSX provides a table where each row corresponds to a protein ortholog group and columns showcase statistics and various information for them. The interactive plot in the HTML output file shows a scatterplot where each point corresponds to a protein ortholog group, the x-axis is the number of genomes the ortholog group is found in, and the y-axis is a metric that indicates syntenic conservation of surrounding genomic contexts. The coloring of the points indicates whether ortholog groups are primarily associated with MGEs (red) or not (grey).

Both reports feature a metric called the context conservation score that is calculated for each ortholog group and represents how conserved their contexts are. This score is the average number of distinct ortholog groups found in the flanking 10 kb genomic contexts of the focal ortholog group divided by the total number of distinct surrounding ortholog groups observed across all its instances. Values closer to zero indicate that context conservation is low, whereas values closer to one indicate the context of the focal ortholog group is highly conserved. The default window size for the extent of surrounding context assessed is 10 kb but this can be adjusted by users via a parameter in bofasa run. The statistic is computed in two ways: 1.) using all instances of the ortholog group and 2.) using only complete instances where surrounding contexts can be properly expanded (i.e. not near scaffold edges).

In addition to the standard output files described, bofasa run also features two options to perform auxiliary analyses and generate additional results. One option, triggered by specifying “-cg” or “--core-genome”, instructs bofasa run to construct MSAs of single-copy, near-core ortholog groups (default requirement of single-core in 95% of genomes, but is adjustable via the “-ns” option). To do so, it assesses domain-resolution ortholog groups that are found in single-copy in 95% of genomes. Note, ortholog groups can be still considered part of the single-copy near-core if they feature multiple proteins in a small subset of genomes. For instance, if an ortholog group is found in 95 of 100 genomes in single-copy, missing in three genomes, and found in two copies in two of the genomes, then, based on default configurations, it will be regarded as part of the single-copy core genome. However, the instances of the ortholog group in the two genomes with multiple copies/paralogs will not be included in the resulting MSA. Domain-resolution ortholog groups are further filtered if they are homologous to insertion sequences^18^ or are predicted to be on plasmids or phages by geNomad^17^. Individual domain ortholog-group specific MSAs are finally concatenated into a single multi-FASTA file. Ultimately, both the directory of multiple sequence alignments for each near-core domain-resolution ortholog groups or the concatenated alignment can be used to perform phylogenomic modeling with software such as FastTree 2 or IQ-TREE. However, given the default behavior to occasionally mask rare instances of likely duplications, we encourage users to perform partition aware phylogenomic modeling. Another argument for bofasa run, “-ogc” or “--og-consensus”, triggers the construction of protein-resolution ortholog group MSAs using MUSCLE^68^ and, subsequently, profile HMMs and consensus sequences using HMMER^72^ (v3.4) to enable efficient identification of established ortholog groups in new genomes. Additional scripts are also provided in the bofasa software suite for format manipulation of result files from bofasa and other orthology prediction software to aid comparisons of orthology predictions.

### Comparison of methods for multi-species orthology prediction

#### Data

All genomes used for showcases are publicly available on NCBI. Genome accessions are provided in **Supplementary Table S1**.

#### Visualization

The ggplot2 library in R was used to visualize scatterplots, barplots, and heatmaps. The iTol server was used to visualize phylogenies^73^. To demonstrate homology between related instances of mobile genetic elements and biosynthetic gene clusters, the software package clinker (v 0.0.31)^37^ and the gggenes library in R were applied.

#### Assessing computational resource requirements

We ran comparisons on a server running Ubuntu 24.04 with 472 Gb memory and 96 threads. Verbose output from the Unix command “time” was used to assess run time to completion and memory consumption, using the “Maximum resident set size” metric as a proxy for memory requirements. For datasets where these performance metrics were assessed, equivalent numbers of threads were specified/requested for each software to make resource comparison fair. For each showcase, details on commands issued for orthology prediction software can be found in the Git repository associated with the manuscript under the sub-directories “Program_Logs/”.

#### SimPan evolutionary simulation

The SimPan workflow^27^ (downloaded on Jan. 31st, 2025), which leverages SimBac^28^ (v0.1a) and INDELible^74^ (v1.03), was used to simulate pangenomes from a common ancestral and artificial bacterial-like genome. We modified parameters to simulate evolutionary divergence that is typically observed between genomes belonging to different species but that are from the same genus. Specifically, SimPan was run using the following options: “--idenOrtholog 0.8 --idenParalog 0.5 --idenDuplication 0.95”. After running SimPan to generate 20 diverse genomes, we reformatted them to GenBank format using the script convertSimPanToGenBankFormat.py, included in the GitHub repository associated with this manuscript. These were then provided as input to bofasa prep to process and the resulting proteomes were primarily used as inputs for various software for orthology prediction. Versions 2.5.5 and 3.1.3 of OrthoFinder were run using the option “-M msa”. SonicParanoid (v2.0.8) was run without Pfam analysis (“--graph-only”) because simulated genomes are not real and their coding sequences do not contain domains. PIRATE (v1.0.5) and SCARAP (v1.0.0) were run using default settings. Finally, bofasa run was executed both with and without the “-rs 1000” parameter which requests for a more comprehensive assessment when partitioning domain-resolution ortholog groups. The scripts, extract_og_pairs.py and compare_og_pairs.py, included within bofasa, were used to list pairs of proteins belonging to the same ortholog groups and compare such lists produced by different methods, respectively.

#### Application to the multi-phylum dataset

A set of nine genomes were selected to represent a small multi-phylum set of genomes to use to assess differences in orthology inference by different methods. Equivalent applications of the programs were performed with and without the inclusion of the *Wolbachia pipientis* genome (GCF_000008025.1). Both versions of OrthoFinder, PIRATE, and SCARAP were run using equivalent options for the SimPan-related showcase. For PIRATE, we additionally ran it using looser cutoffs for determining orthology by specifying the option: “ --steps 20,30,40,50,60,70,80,90,95,98”. SonicParanoid was run using both default parameters and the option “-m sensitive” to reflect the diversity of the genome set. Similarly, bofasa was executed with default parameters as well as with the prep option “-emg” and run options “-rs 1000 -us”. KOfam profile HMMs were used to annotate proteins to predefined KEGG ortholog groups with KofamScan (v1.3.0)^26^. Comparisons between methods were performed using the scripts extract_og_pairs.py and extract_scc_og_pairs.py to assess the total number of ortholog pairs and the size of the strict single-copy core genome. The script determine_og_freqs.py was used to determine ortholog group sizes and assess differences in the cumulative distributions of proteins clustered by different methods when ranked in order of smallest to largest ortholog groups.

#### Comparative analysis of the Staphylococcus genus

A representative genome was selected for each staphylococcal species in GTDB R226 which had a complete-level genome available^3^. Both versions of OrthoFinder, PIRATE, SCARAP, and SonicParanoid were run using default options. Two bofasa executions were performed using default options as well as the “-emg” prep option together with the run option “-rs 1000”. Comparisons between methods were performed using the scripts extract_og_pairs.py, extract_scc_og_pairs.py, and determine_og_freqs.py.

To assess differences in the size of the core genome, we generated MSAs for strict single-copy core genes using MUSCLE super5 and combined them prior to trimming for sites that were largely ambiguous, ≥10% of sites were “-” or “X”. Afterwards, the length and the total percentage of positions across all genomes that correspond to gaps were computed for each core genome inferred by a method. This was also performed the same way for both bofasa runs; however, we also reran bofasa using default options with the “-cg” flag to showcase its improved extraction of a large core genome that is also free of sequences associated with mobile-genetic elements, which likely do not represent vertical evolution, and inter-domain regions, which are likely more affected by genetic drift than selection.

To investigate the distribution of integrases, we annotated ortholog groups for their presence using associated domains according to the authors of TIGER^75^, note some corresponded to housekeeping proteins such as the recombinase XerC involved in DNA replication. These were similarly used to identify ortholog groups that corresponded to or are related to integrases by other orthology prediction software. Some proteins belonging to ortholog groups that featured proteins with integrase associated domains did not feature such domains themselves and were regarded as “integrases by association”. The functions of such proteins were further assessed using structure prediction via both AlphaFold^36^ and Phyre 2^35^.

We developed the program contextCrawler.py to compute the standardized context conservation score for every inner-node of the phylogeny for the largest integrase-associated ortholog group, OG01033, available in the repository associated with the bofasa manuscript (https://github.com/Kalan-Lab/bofasa_manuscript/blob/main/showcases/staph_genus/integrase_investigation/Fig4c/contextCrawler.py). The standardized variant of the context conservation score was computed as the context conservation score for the protein children of the inner-node divided by the median context conservation score observed across ortholog groups of the same size. The program fai in the zol suite (v1.6.19) was used to search for homologous instances of the *i2* integrase from OG01033 via single-query mode. In order to identify a closely related *Staphylococcus aureus* genome to one that harbors the *i2* integrase, skani (v0.3.2) was used to compare it to other *Staphylococcus aureus* genomes in GTDB R226. The MGE *mini-MGE 14 (mM14)* was delineated manually based on region-specific alignment between the query genome (GCA_900620215.1) and its nearest neighbor without the integrase (GCA_947040485.1). To identify additional instances of MGEs related to *mM14*, we subsequently used it as a query to remotely search NCBI’s nr database with cblaster (v1.4.0).

#### Investigating multi-domain biosynthetic genes across the Actinomycetota phylum

We similarly ran multiple methods to compare their performance against bofasa on a set of 56 Actinomycetota genomes. These genomes were representative of distinct genera in the phylum from GTDB release 220 which had large genomes >5 Mb and were designated as complete. A species tree for the genomes was constructed using GToTree (v1.8.8) with midpoint rooting applied.

The software antiSMASH^47^ (v7.0.0) was used to annotate biosynthetic gene clusters across genomes and identify NRPSs. Product predictions in antiSMASH results were assessed to determine whether NRPS-containing BGC regions encoded for metallophores. To manually assess the contexts of select NRPS-related ortholog groups highlighted in the manuscript for metallophore biosynthesis^48^ and transport genes^49^, we used HMM profiles provided as supplementary materials from Reitz et al. 2026. Standardized scores for enzyme profiles were computed as the maximum score observed in the flanking contexts of focal NRPS genes divided by the threshold score needed for their detection.

#### Application to MAG-based showcases

Relative abundance information for metagenome assembled genomes (MAGs) from Yaffe et al. 2025 were assessed to identify the top hundred bacterial species in healthy microbiomes. Specifically, only abundances for timepoints A1-A4, reflective of microbiomes prior to shifts due to self-administered antibiotics, were used to profile the microbiomes of participants in the study who had largely reported as being healthy. The median relative abundance was calculated for each MAG across these timepoints for all participants (with abundances of zero included) and ranked to determine the top hundred most-abundant species within gut microbiomes that also had complete genome assemblies available in GTDB release 226.

Complete genome assemblies for the top hundred species were processed through bofasa prep and used to perform orthology inference with different methods. To assess how assembly quality impacts orthology prediction, we further developed the program gappy (v1.0.1; https://github.com/raufs/gappy) to simulate assembly fragmentation and base-calling errors. The gappy code was largely developed using AI with subsequent manual editing and vetting. Briefly, it samples gap-lengths from a beta distribution (default parameters: α=1.09, β=10.0, range=[1, 1M bp]) and randomly introduces gaps into assemblies. The algorithm terminates when a desired percentage of the input genome assembly, for instance 75%, is retained. For demonstrations in this manuscript, we requested assembly fragmentation to retain 50% and 75% of the input complete-level genome assemblies. After generating these artificially fragmented genomes, we processed them through bofasa prep and reperformed orthology inference using different methods. To assess the impacts of fragmentation, we mapped coding gene annotations for fragmented genomes to their respective versions in the complete-level genomes.

## Notes

https://github.com/raufs/bofasa

### Competing Interest Statement

The authors have declared no competing interest.

