## Supplementary Figures for "*bofasa* enables high-quality orthology inference across diverse multi-species bacterial genome sets"

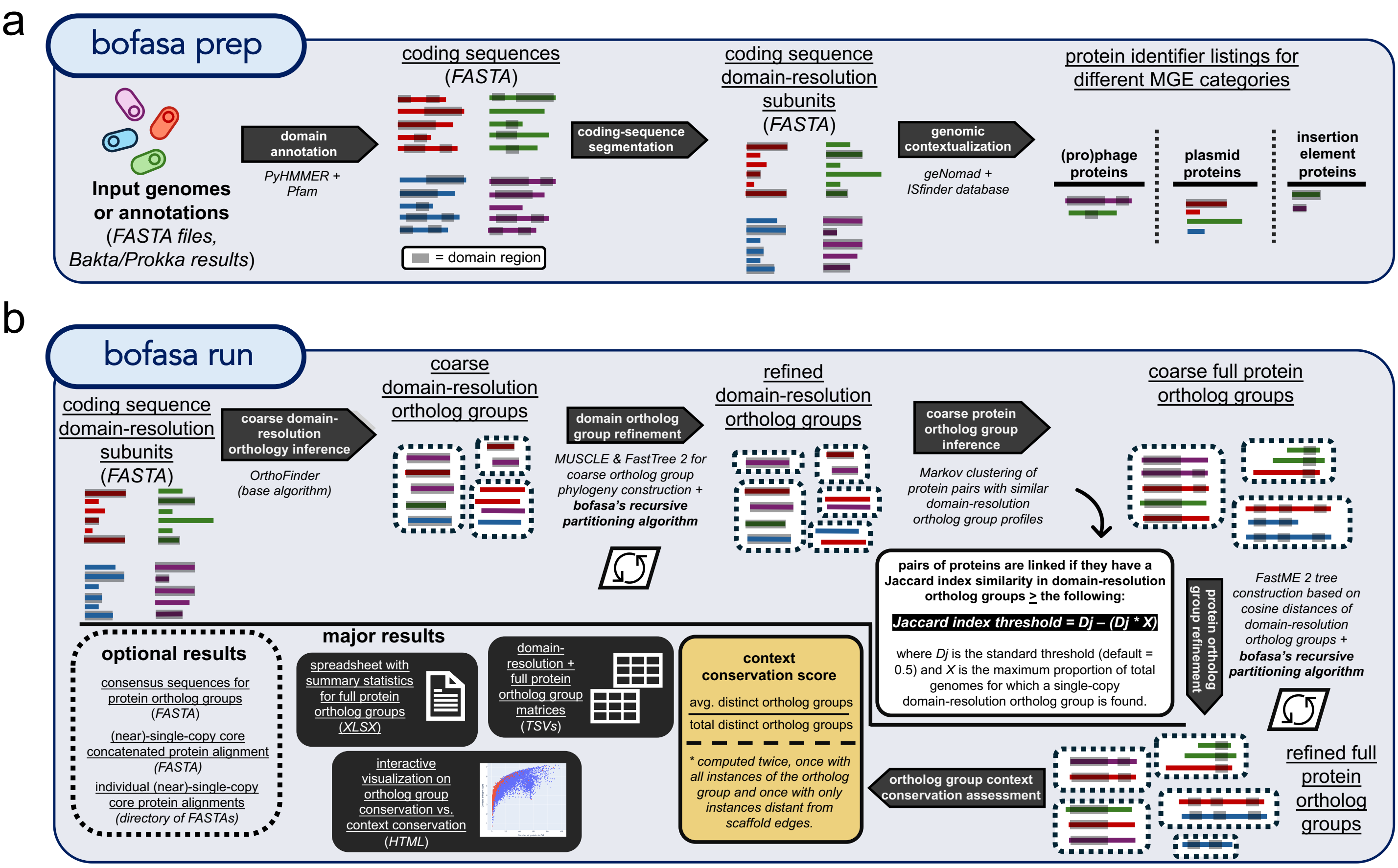


**Supplementary Figure 1: Overview of the bofasa toolkit.** A graphical overview is provided for the core framework of (**a**) bofasa prep and (**b**) bofasa run. For an overview of bofasa's recursive partitioning algorithm, see **Fig. 2**.

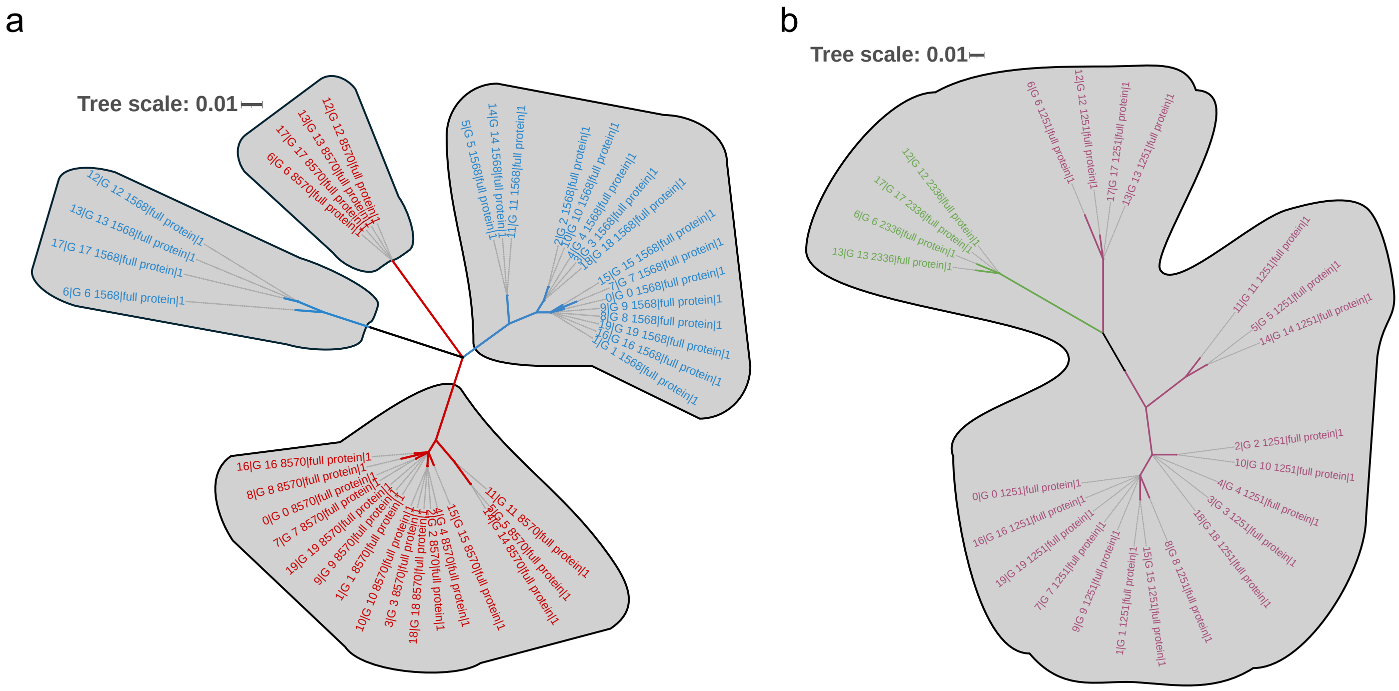


**Supplementary Figure 2: Examples of false positive and negative orthology prediction by bofasa.** (**a**) An example of missed orthology predictions (false negative; under-clustering) by bofasa. (**b**) An example of false positive orthology predictions (over-clustering) by bofasa. In both cases, approximate maximum likelihood phylogenies for coarse domain-resolution ortholog groups are constructed from protein alignments. The color of the leaves represents actual ancestral origins of genes based on tracing from simulation. The grey outlines of specific phylogenetic clades represent the delineations of resolved ortholog groups by bofasa.

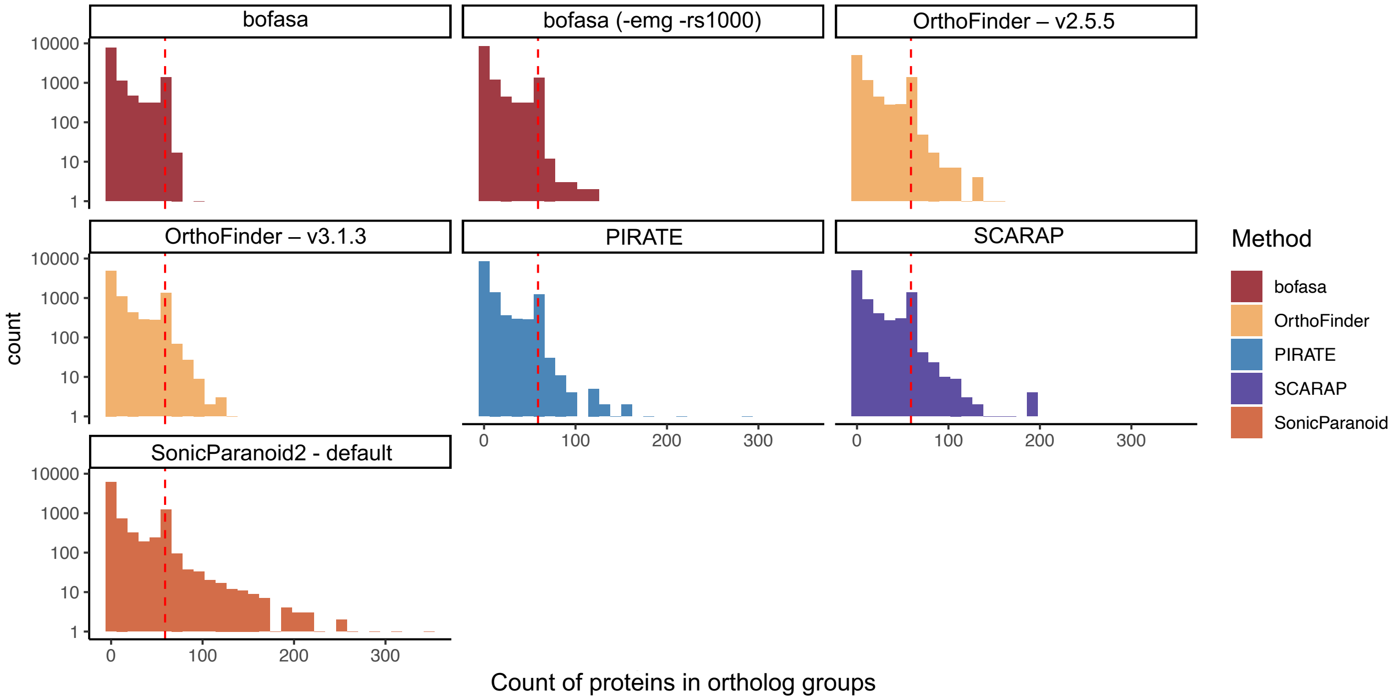


**Supplemental Figure 3: Ortholog group size distributions vary across prediction software.** Histograms of the number of proteins found within ortholog groups from application of different orthology prediction software on the dataset of 59 complete genomes representative of distinct *Staphylococcus* species.

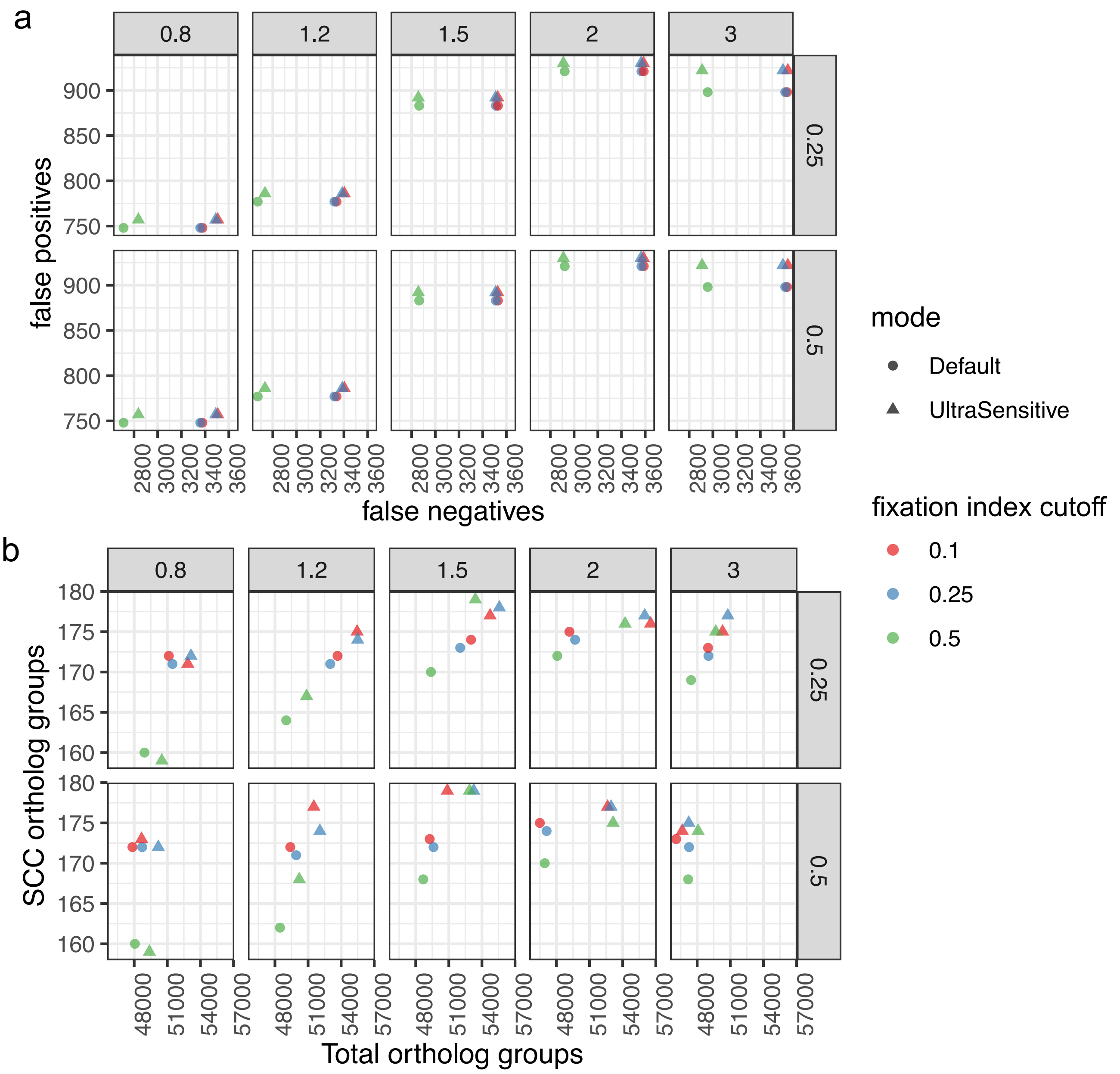


**Supplementary Figure 4: Parameter optimization for different applications.** The impacts of different parameter values are assessed as combinations on the (**a**) SimPan and (**b**) multi-phylum genomic datasets. (**a**) For the SimPan dataset, the number of false negatives and false positives is shown, with parameter combinations lowering both metrics preferred. (**b**) For the multi-phylum dataset, the number of single-copy core (SCC) ortholog groups identified and the total number of pairs of proteins belonging to the same ortholog group were assessed. Preference was given to parameter combinations which increased the number of SCC ortholog groups while reducing the number of total orthologs inferred. Grids are arranged vertically by MCL inflation values provided to OrthoFinder for determination of coarse domain ortholog groups. They are arranged horizontally by Jaccard index thresholds for shared resolute domain ortholog groups needed to identify pairs of full proteins as homologous, used in the process of determining full protein ortholog groups. The color of runs corresponds to different values for the fixation index cutoff, which controls re-merging of resolute domain ortholog groups. Their shape indicates whether OrthoFinder was run using the ultra-sensitive DIAMOND search flag or not.

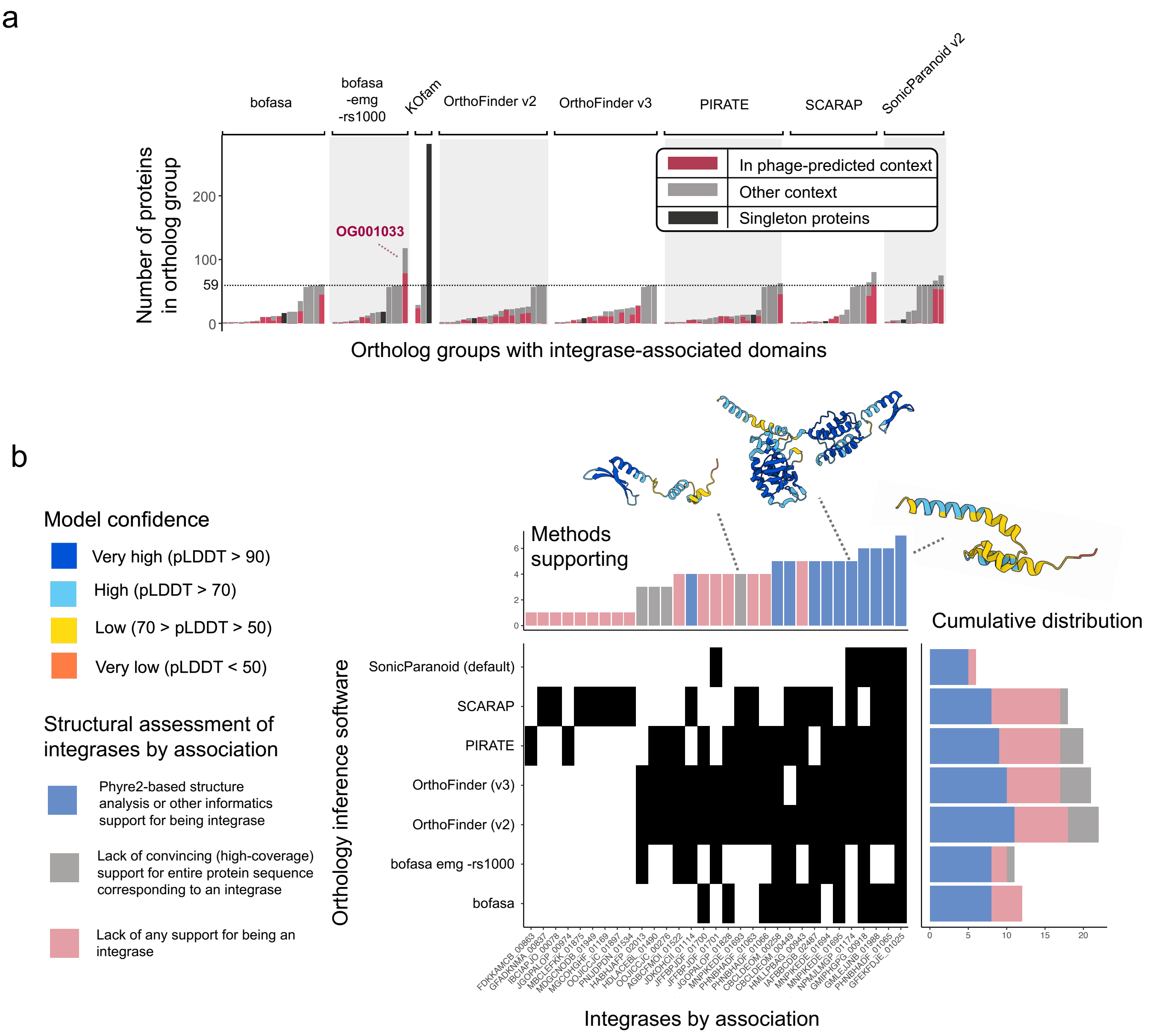


**Supplemental Figure 5: Comparisons of integrase-associated ortholog group prediction by different methods.** (**a**) The size of ortholog groups encoding for one or more proteins with integrase-associated domains is shown across different methods. The coloring corresponds to the proportion of proteins within each ortholog group that were predicted via geNomad to lie on phages. One bar per method, shown in black, correspond to the number of singleton proteins, not orthologous to other proteins. (**b**) The identification of proteins regarded as "integrases by association" (lacking the associated domains themselves but grouped with other such proteins) is shown. Structural information was used to further assess whether such proteins were likely actually integrases or not and to assess differences between orthology prediction methods.

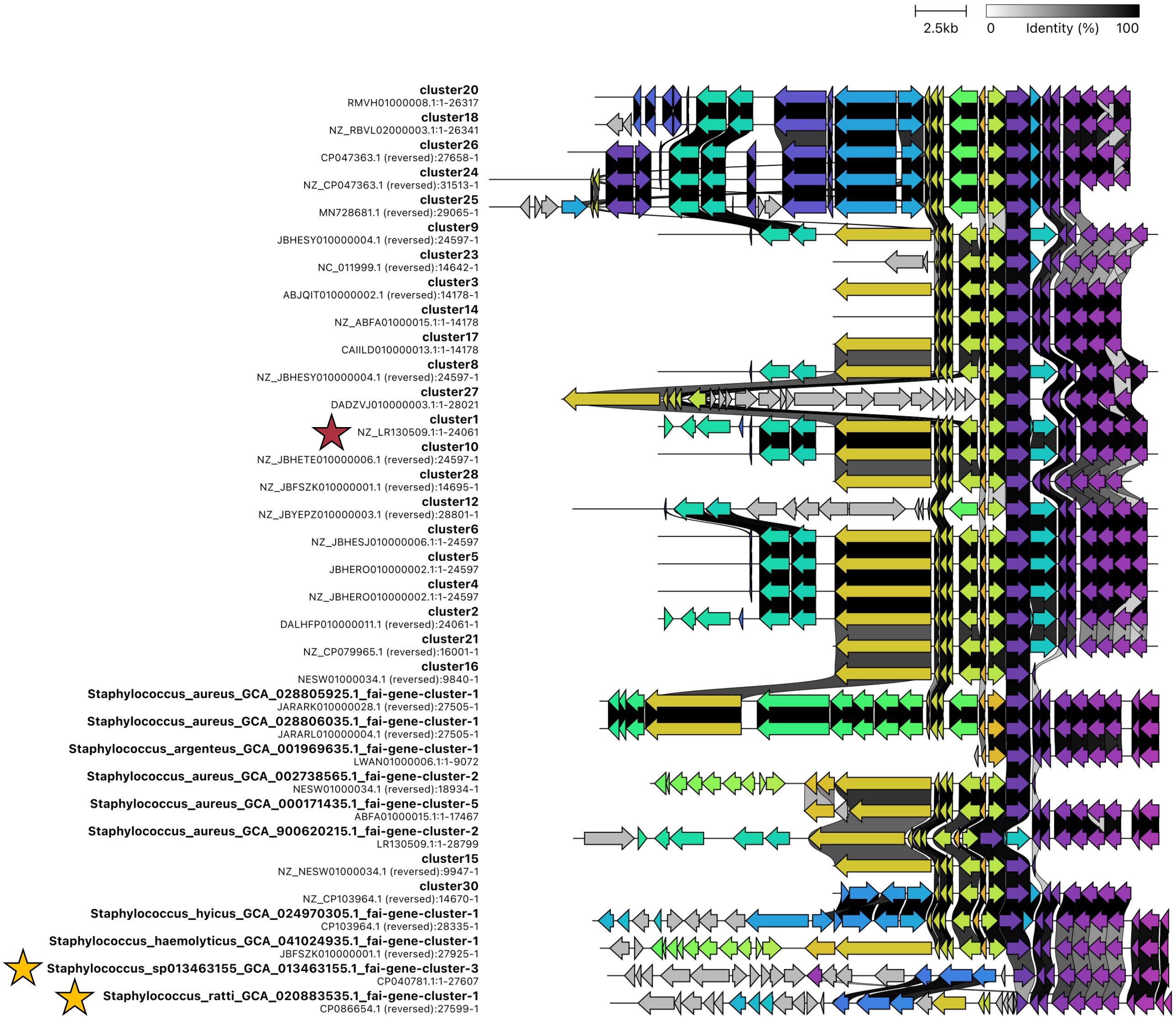


**Supplemental Figure 6: Comparisons of genomic regions containing *i2*-homologs and homologous instances of *mM14*.** A visualization created using clinker of the homology between genomic regions either corresponding to those found via fai-based analysis as containing highly homologous instances of the *i2* integrase from OG01033 or via cblaster based searching of *mM14* in NCBI's nr database. The red star indicates *mM14* and the two gold stars correspond to the two genomic neighborhoods shown in **Fig. 4d** that contain the *i1* and *i2* integrases from OG01033.

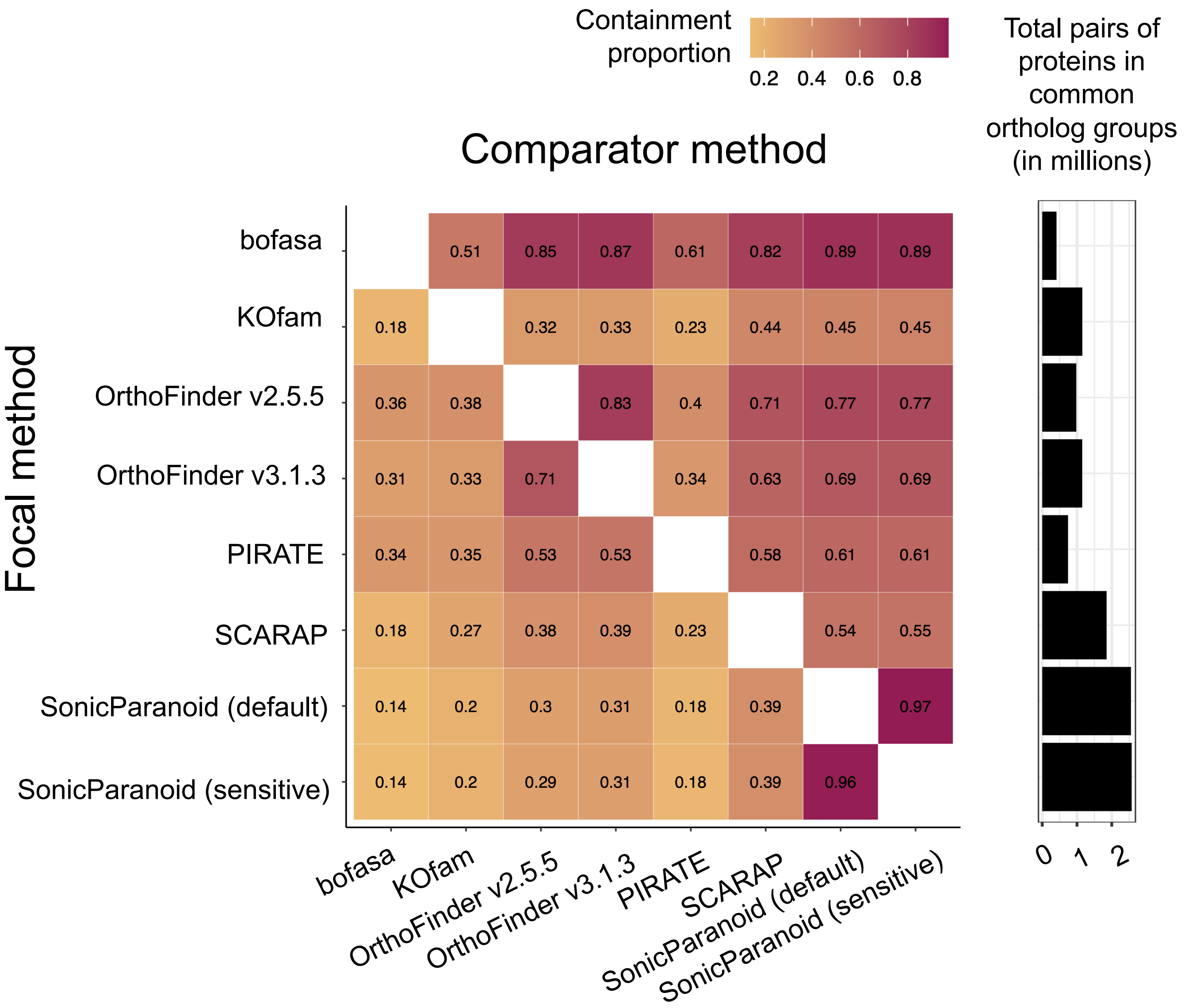


**Supplementary Figure 7: Comparisons of different orthology prediction software on 56 Actinomycetota genomes.** A heatmap showing the proportion of ortholog pairs for focal methods (rows) contained by comparator methods (columns). A barchart of the total number of ortholog pairs predicted by each method is shown on the right.

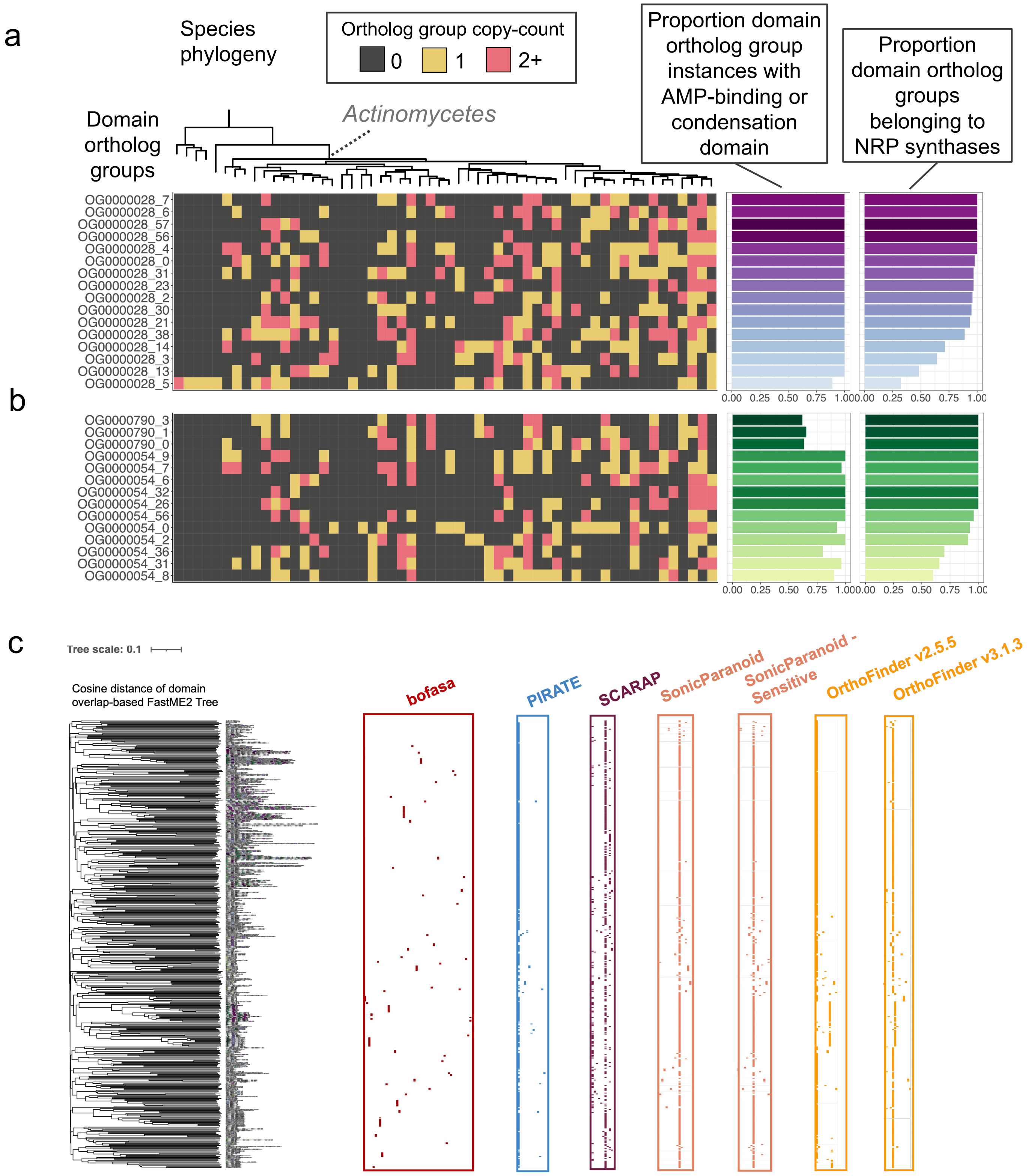


**Supplementary Figure 8: Assessing NRPS-associated orthology predictions by bofasa and other methods.** Domain-resolution ortholog groups detected by bofasa where at least 25% of member protein-chunks were found to belong to NRP synthetases annotated by antiSMASH and 50% corresponded to (**a**) an AMP-binding domain or (**b**) condensation domain. The species tree was constructed using GToTree and midpoint-rooted. Coloration of the heatmap indicates the copy-count of domain-resolution ortholog groups across the 59 Actinomycetota genomes. (**c**) A tree based on domain compositions is shown for full NRP synthetase proteins from the 59 Actinomycetota genomes as depicted in Fig. 5c. Distributions of protein-resolution ortholog groups that include two or more NRP synthetases are shown for different orthology prediction software.

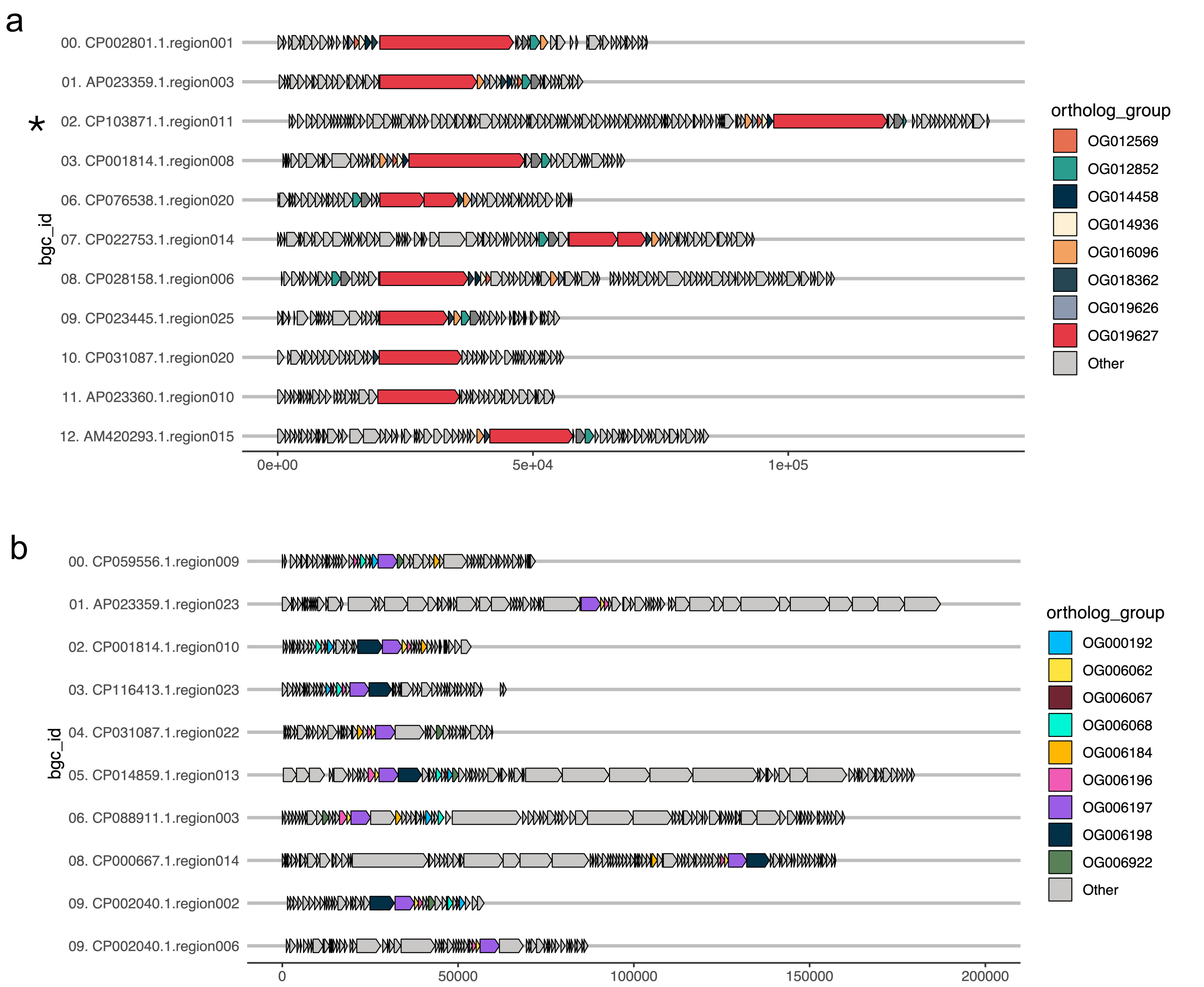


**Supplementary Figure 9: Assessing the contexts of the two largest NRPS-associated ortholog groups identified by bofasa across Actinomycetota genomes.** The contexts of proteins belonging to the NRPS-associated ortholog groups (**a**) OG019627 [red] and (**b**) OG006197 [purple] highlighted in **Fig 5cde** are shown. The contexts represent BGC regions predicted by antiSMASH with the focal ortholog groups. Common ortholog groups found in five or more BGC regions are distinctly colored. * The BGC region was restricted to the first 200,000 bp to allow easier visual assessment of the panel.

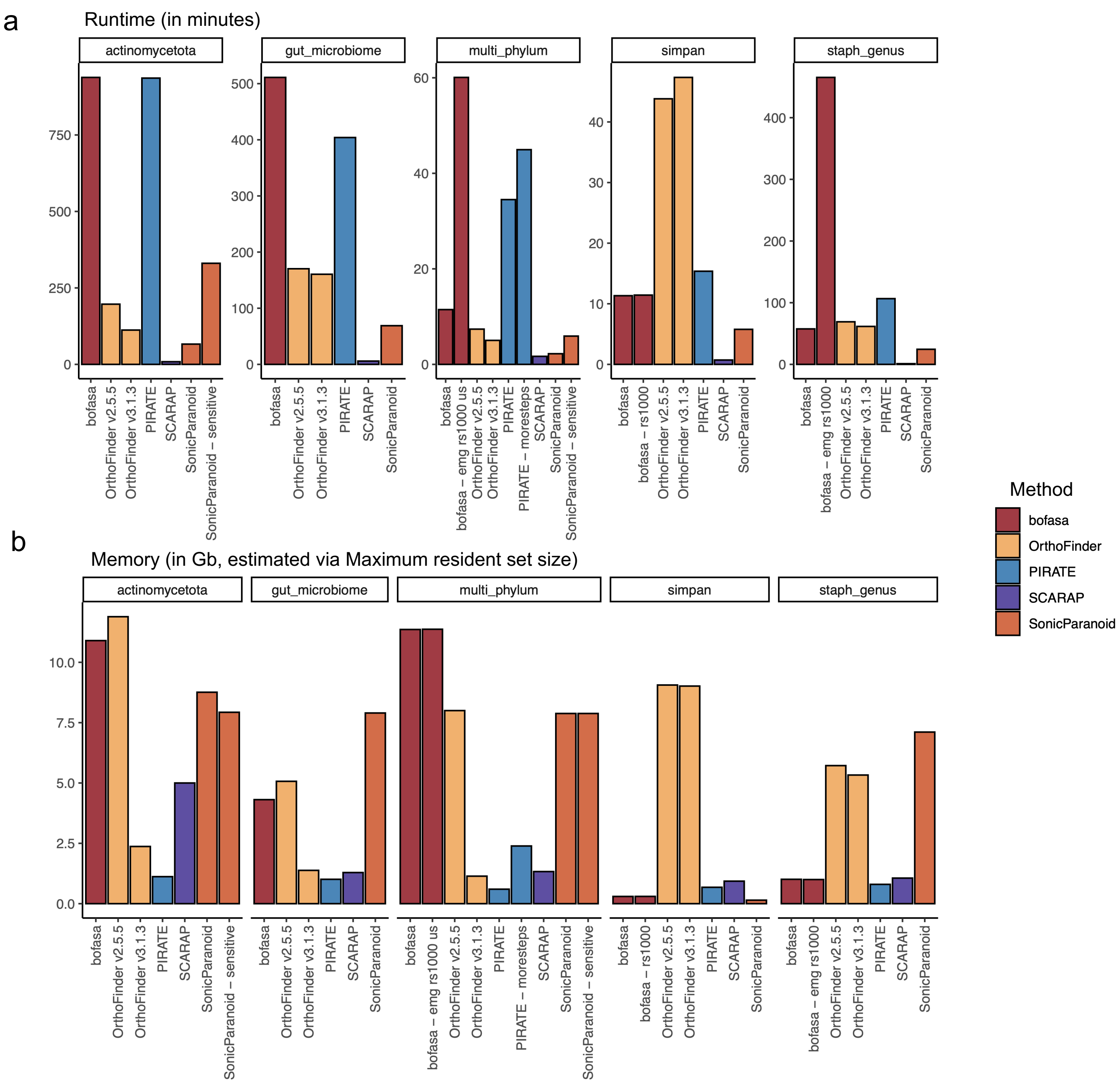


**Supplemental Figure 10: Runtime and memory requirements for orthology prediction software.** The (**a**) runtime and (**b**) memory, proxied via the "Maximum resident set size" statistic reported by the unix command *time,* are shown for different orthology prediction methods across the five showcases in the manuscript.
